# The structure of a hibernating ribosome in a Lyme disease pathogen

**DOI:** 10.1101/2023.04.16.537070

**Authors:** Manjuli R. Sharma, Swati R. Manjari, Ekansh K. Agrawal, Pooja Keshavan, Ravi K. Koripella, Soneya Majumdar, Ashley L. Marcinkiewicz, Yi-Pin Lin, Rajendra K. Agrawal, Nilesh K. Banavali

**Author notes:** University of California at Berkeley, Berkeley, CA.

## Abstract

The spirochete bacterial pathogen *Borrelia* (*Borreliella) burgdorferi* (*Bbu*) affects more than 10% of the world population and causes Lyme disease in about half a million people in the US annually. Therapy for Lyme disease includes antibiotics that target the *Bbu* ribosome. We determined the structure of the *Bbu* 70S ribosome by single particle cryo-electron microscopy (cryo-EM) at a resolution of 2.9 Å, revealing its distinctive features. In contrast to a previous study suggesting that the single hibernation promoting factor protein present in *Bbu* (bbHPF) may not bind to its ribosome, our structure reveals a clear density for bbHPF bound to the decoding center of the small ribosomal 30S subunit. The 30S subunit has a non-annotated ribosomal protein, bS22, that has been found only in mycobacteria and Bacteroidetes so far. The protein bL38, recently discovered in Bacteroidetes, is also present in the *Bbu* large 50S ribosomal subunit. The protein bL37, previously seen only in mycobacterial ribosomes, is replaced by an N-terminal α-helical extension of uL30, suggesting that the two bacterial ribosomal proteins uL30 and bL37 may have evolved from one longer uL30 protein. The longer uL30 protein interacts with both the 23S rRNA and the 5S rRNA, is near the peptidyl transferase center (PTC), and could impart greater stability to this region. Its analogy to proteins uL30m and mL63 in mammalian mitochondrial ribosomes also suggests a plausible evolutionary pathway for expansion of protein content in mammalian mitochondrial ribosomes. Computational binding free energies are predicted for antibiotics, bound to the decoding center or PTC and are in clinical use for Lyme disease, that account for subtle distinctions in antibiotic-binding regions in the *Bbu* ribosome structure. Besides revealing unanticipated structural and compositional features for the *Bbu* ribosome, our study thus provides groundwork to enable ribosome-targeted antibiotic design for more effective treatment of Lyme disease.

---

Lyme disease affects up to 14.5 % of the human population worldwide^1^ and is the most prevalent tick-borne disease in the Northern hemisphere, including the United States^2^. Its causative agent is the spirochete bacteria genospecies complex, *Borrelia burgdorferi sensu lato* (also known as *Borreliella burgdorferi sensu lato* or Lyme borreliae), which includes the species *Borrelia burgdorferi sensu stricto* (*Bbu*), the primary cause of human Lyme disease in North America. Elongated seasons and widened habitats for the tick vectors driven by climate change are resulting in its continued rise^3^. *Bbu* transmitted from tick saliva into human skin after the tick bite can spread and affect the human cardiovascular and nervous systems^4^. The direct economic burden of Lyme disease on the United States healthcare system alone is estimated to be a multi-billion amount each year^5^. The ribosome is an RNA-protein molecular machine that coordinates the vital process of protein synthesis in all living organisms^6^. Over two decades of detailed structural studies on various prokaryotic and eukaryotic ribosomes have clarified various aspects of the four mechanistic steps of translation – initiation, elongation, termination, and recycling^7-9^. However, even in prokaryotes, new 70S ribosome structures continue to reveal unexpected functional and compositional features, such as the discovery of structural basis of specific ribosome hibernation mechanisms^10-17^ and presence of new smaller ribosomal proteins^14,18-20^. When diagnosed early in the infection, Lyme infections can be adequately treated with antibiotics, including ribosome-targeting antibiotics such as doxycycline and erythromycin^21^. Recently, hygromycin A (HygA), a ribosomal large (50S) subunit-binding antibiotic, was discovered to be extremely selective in resolving *Bbu* infections^22^. Knowledge of the structural details of the *Bbu* ribosome are therefore highly relevant for designing better biomedical interventions for Lyme disease.

In this study, we report the 70S *Bbu* ribosome structure in its hibernating state solved at a resolution of 2.9 Å using single particle cryogenic electron microscopy (cryo-EM). The local resolution of the *Bbu* ribosome density is near the Nyquist limit of 2.2 Å in the core regions of the ribosome with the flexible regions such as parts of the small (30S) subunit head, the 50S subunit components such as uL1 stalk, the uL7/uL12-stalk base and uL9 showing lower resolution (Figure 1A, supplementary movie 1). The structure contains 58 resolved components (Table S1, Figure 1B): 3 ribosomal RNAs (23S, 5S, 16S), 1 tRNA in the E-site (Figure 1C, 1D), 21 proteins in the small subunit (uS3 through bS22), and 33 proteins in the large subunit (uL1 through bL38) (Figure S1). Of these, the existence of two previously unknown *Bbu* ribosomal proteins is revealed directly through their cryo-EM density: bS22 in the small subunit (Figure 1C, 1E) and bL38 in the large subunit (Figure 1C, 1F). The uL30 protein in *Bbu* is enlarged through an N-terminal extension (Figure 1C, 1G) that occupies the same site as the much smaller helical ribosomal protein bL37 in mycobacterial ribosomes.

**Figure 1.**
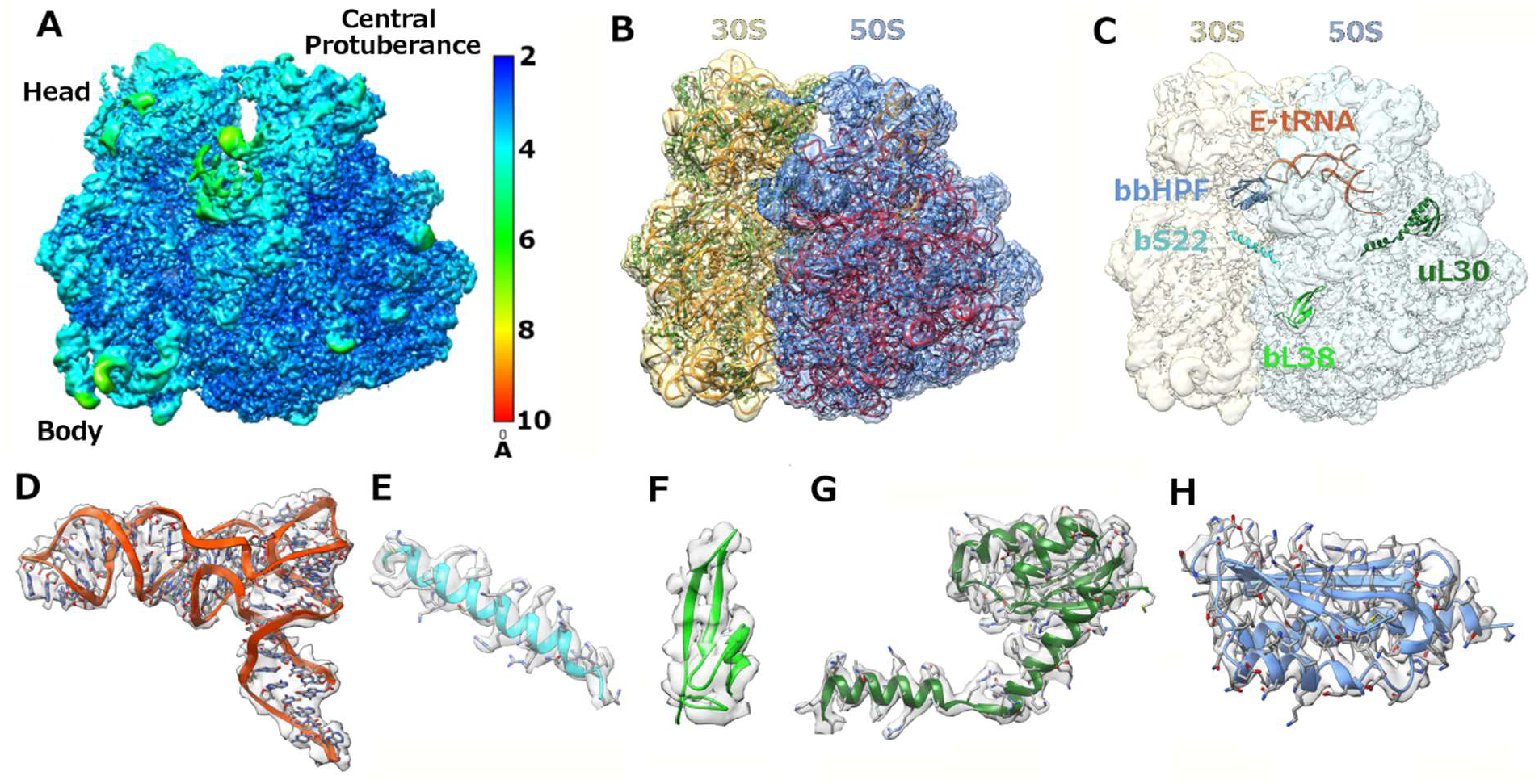
*Bbu* 70S hibernating ribosome structure and protein components. **(A)** Cryo-EM density map of 70S with local resolution indicated by color. **(B)** Model fit into cryo-EM density map with 30S density in khaki, 50S density in blue, 16S RNA and 5S RNA in orange, 23S RNA in red, 30S proteins in green, and 50S proteins in blue. **(C)** Notable components in the structure shown within the cryo-EM density map. These components of the hibernating 70S ribosome, shown in ribbon format with known sidechains in stick format, are **(D)** the E-site tRNA (orange), **(E)** the bS22 protein (cyan), **(F)** the putative bL38 protein (green), **(G)** the longer uL30 protein (dark green),and **(E)** the bbHPF protein (blue).

The single 97 amino-acid (aa) residue hibernation promoting factor (HPF) gene in *Bbu*, named BB_0449 in the KEGG database^23^ (Uniprot ID: A0A8F9U6W1), was previously reported to have low mRNA transcript and expressed protein levels during various growth phases^23^. The expressed protein did not localize to the ribosome-associated protein fraction, and its gene deletion did not affect the mouse-tick infectious cycle^24^. Based on these observations, it was suggested that the 97 aa-residue *Bbu* HPF protein (bbHPF) may not play its traditional role in ribosome hibernation^24^. The bbHPF protein is a “short” HPF resembling the *E. coli* YfiA and HPF proteins and differs from “long” HPFs in lacking a C-terminal dimerization domain, which forms a dimerization interface that stabilizes a hibernating 100S ribosome dimer in certain bacteria^10,12,15,17^. In our structure, we find bbHPF bound to the decoding center in the small subunit of the *Bbu* 70S monosome, thereby confirming the sequence-based expectation that it does play a role in 70S ribosome hibernation (Figure 1C, 1H). An E-site tRNA was found interacting with the C-terminal end of bbHPF with no mRNA density near it. Maintenance of the bbHPF and E-site tRNA densities in all sub-classes of the 70S ribosome obtained through multiple 3D classifications suggest that bbHPF and E-site tRNA colocalize on the hibernating *Bbu* 70S ribosome and are not a combined density due to two separate HPF-bound or E-site tRNA-bound *Bbu* 70S ribosome populations.

Including the bbHPF structure reported in this work, atomic resolution structures for eight HPF proteins from seven different bacterial species are now known (Table S2). When aligned by primary protein sequence, multiple positions in these HPF proteins have similar amino acids, but Leu83 is only fully conserved amino acid residue (Tables 1 and S3). This suggests that a certain level of sequence divergence is tolerated in the HPF domain that binds at the decoding center of bacterial ribosomes during their hibernation. This HPF domain is similar in all these bacterial species, having a fold with two α-helices and four β-strands in a β1-α1-β2-β3-β4-α2 topology, with β1 and β2 strands forming a parallel β-sheet and the β2, β3, and β4 strands forming an antiparallel β-sheet continuous with the first parallel β-sheet. There seems to be permissibility for an increase in the size of some of the β strands or α helices (e.g. β1 or α1), a break in the helicity of an α-helix (α1), and increases in the sizes of the loops between β-strands (β2-β3 or β3-β4) (Table 1). The YfiA protein (Uniprot ID: P0AD49) and the HPF protein (Uniprot ID: P0AFX0) in *Escherichia coli* (*Eco*) differ in sequence but both bind the same decoding center site in the *Eco* ribosome, indicating intra-species sequence permissibility in HPF-ribosome interaction. There are multiple structures of the *Eco* YfiA bound to the *Thermus thermophilus* (*Tth*) ribosome^16,25-31^, indicating that inter-species permissibility also exists in HPF-ribosome interaction. There is also internal structural variability in the HPFs, as seen in the overlay of these known HPF structures aligned to each other (Figure 2B), and additional positional variability of the HPFs in their binding pocket, as seen in the overlay of these known HPF structures bound to a bacterial ribosome with the structures aligned using the 16S ribosomal RNA (Figure 2C). Taken together, this evidence suggests that occupation of the bacterial small subunit decoding center binding site does not require a highly conserved HPF sequence, a very rigid HPF structure, or a very restrictive positioning of the HPF in its ribosomal binding site.

**Table 1.**
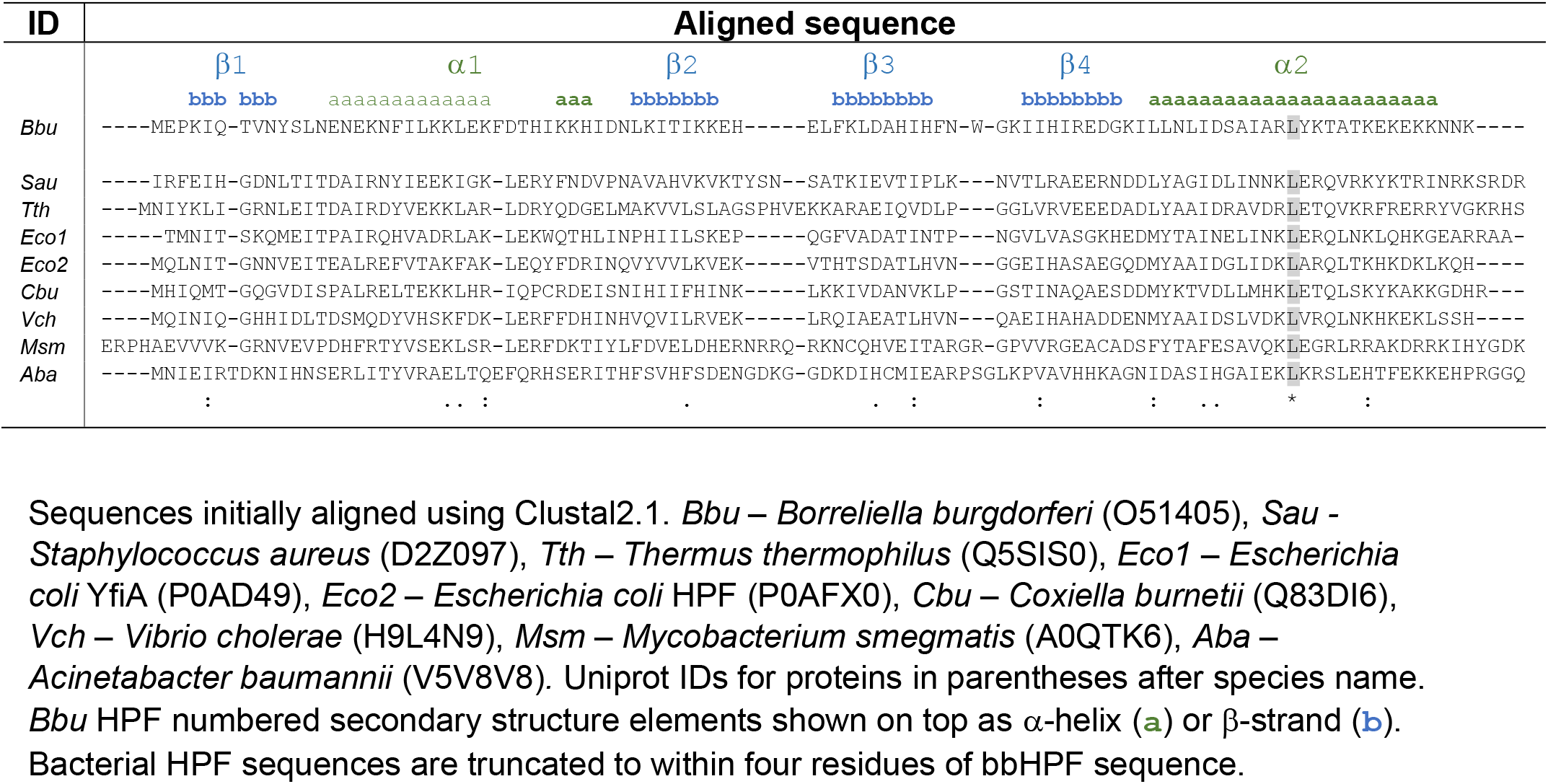
Curated sequence alignment for bacterial HPFs with known structures.

**Figure 2.**
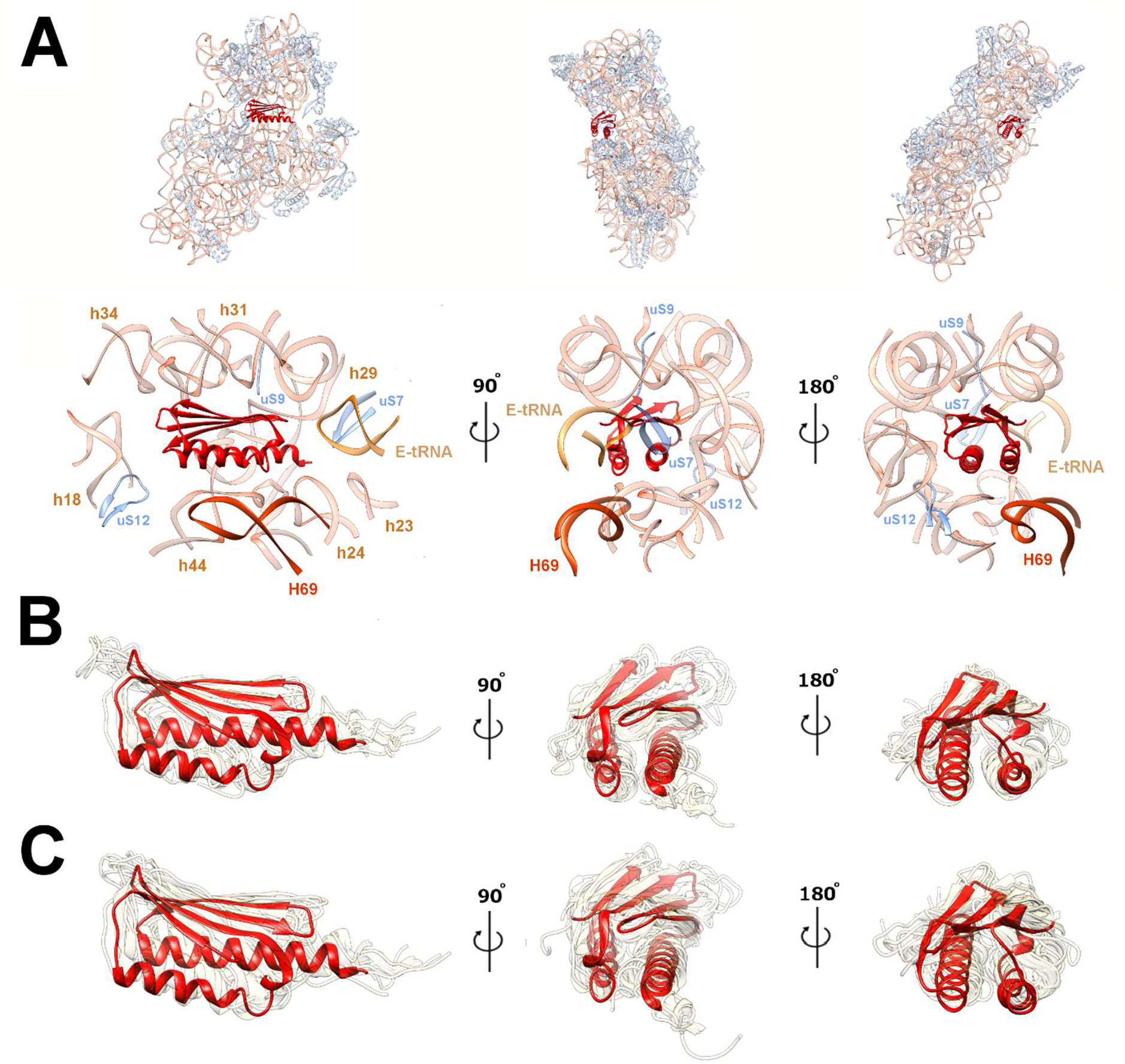
The bbHPF binding site orientation and bacterial HPF protein structural variability. **(A)** Subunit interface (left), mRNA exit (middle) and mRNA entry (right) site views of bbHPF in its decoding center binding site in the *Bbu* ribosome with full 30S view shown above. Labels indicate location of ribosomal proteins (in blue) uS7, uS9, uS12, E-site tRNA (in yellowish orange, labeled E-tRNA), and large subunit RNA helix 69 (in dark orange, labeled H69) of the 23S ribosomal RNA. 16S ribosomal RNA shown in semitransparent light orange. Known bacterial HPF structures, with resolutions better than 3.5 Å, shown in transparent khaki, aligned by HPF protein residues, showing internal structural variability, and **(C)** aligned by 16S ribosomal RNA residues, showing position variability in their ribosomal binding site. BbHPF is shown as opaque red in all panels. Views shown in panels B-C are analogous to those shown in panel A. Details of structures used in panels B-C are listed in Supplementary Table S2.

Including the structure reported in this work, there are four ribosome structures with HPF proteins and E-site tRNA both modeled into the density map. These include a hibernating *Mycobacterium smegmatis* (*Msm*) ribosome structure (PDB ID: 5ZEP^32^) and two *Escherichia coli* (*Eco*) hibernating ribosome structures (PDB IDs: 6H4N^11^, 6Y69^33^). When aligned using the HPF protein, there is a clear relative motion revealed between the HPF protein and the E-tRNA in the four structures (supplementary movie 2). The *Msm* structure and the *Eco* structures show substantially different relative HPF and E-tRNA orientations and the *Bbu* structure shows an intermediate relative orientation (Figure S2). When aligned using the small subunit 16S RNA, both the HPF protein and the E-tRNA adjust their relative positions with respect to the ribosome such that overall binding of the two appears roughly similar in all four structures (Figure S3). There is no mRNA density modeled near the E-tRNA in any of these structures, suggesting that the C-terminal end of the HPF protein and the 30S environment in that region enable the binding of the anti-codon stem loop part of the E-tRNA in hibernating ribosomes.

The *Bbu* small subunit unexpectedly showed a clear density for a helical protein in the binding pocket seen to be occupied by the bS22 protein in mycobacterial^14,18,19^ and Bacteroidetes^20^ ribosome structures. There was no annotated bS22 protein in the *Bbu* genome necessitating identification of the protein sequence through a translated nucleotide genome search. *Borrelia* species can be phylogenetically grouped into three lineages, Lyme borreliae, relapsing fever borreliae, and reptile- and echidna-associated Borrelia^34^. Using an approach described in the Supplementary material section 1.1, the bS22 protein sequence was identified as using a non-canonical GUG start codon and confirmed in two ways: (a) modeling of the protein sequence into the cryo-EM density map and finding good sidechain density fits (Figure 1E), (b) finding the same conserved sequence protein in all three groups of *Borrelia* species (Table S4). The *Bbu* bS22 protein has greater structural and sequence homology with the *Fjo* bS22 protein than the *Msm* bS22 protein (Figure 3, Supporting Information section 1.1). It is highly basic, with 61% of its residues being either lysine or arginine. This presumably aids its binding to an RNA pocket near 16S RNA helices h2, h27, h44, h45, and 23S RNA helix H68 (Figure 3). Amongst the three groups of *Borrelia* species, only one residue change is observed in this protein sequence (K10Q) while all other amino acids are completely conserved (Table S4). This protein is likely to be discovered in other bacterial species since its shorter length and non-canonical start codon possibility clearly pose difficulty for automated protein annotation tools and the ribosomal RNA pocket to which it binds is structurally conserved. Just like *Msm* and *Fjo* bS22 proteins, the *Bbu* bS22 protein is analogous to the mammalian mitochondrial small subunit protein mS38^35-37^ and the mammalian cytosolic protein eL41^38^ (Figure 3E – 3H). The mS38 protein, also called the Aurora kinase A interacting protein, is a longer protein with a bent helical structure in which the bend is at the same location as in the *Bbu, Fjo*, and *Msm* bS22 proteins (Figure 3G). The eL41 protein, although named as a large subunit protein, is localized within the small subunit binding pocket in eukaryotic ribosomes^38^, except for one, where it is also found in a large subunit pocket^39^. It has only one helix with 24 aa residues, is shorter than the *Bbu* bS22 protein, and has no bend in its structure (Figure 3F). These similarities among bS22 and its eukaryotic analogs suggest that these ribosomal proteins qualify for a universal name, instead of them being named separately as distinct bacterial, mitochondrial, or eukaryotic ribosomal proteins.

**Figure 3.**
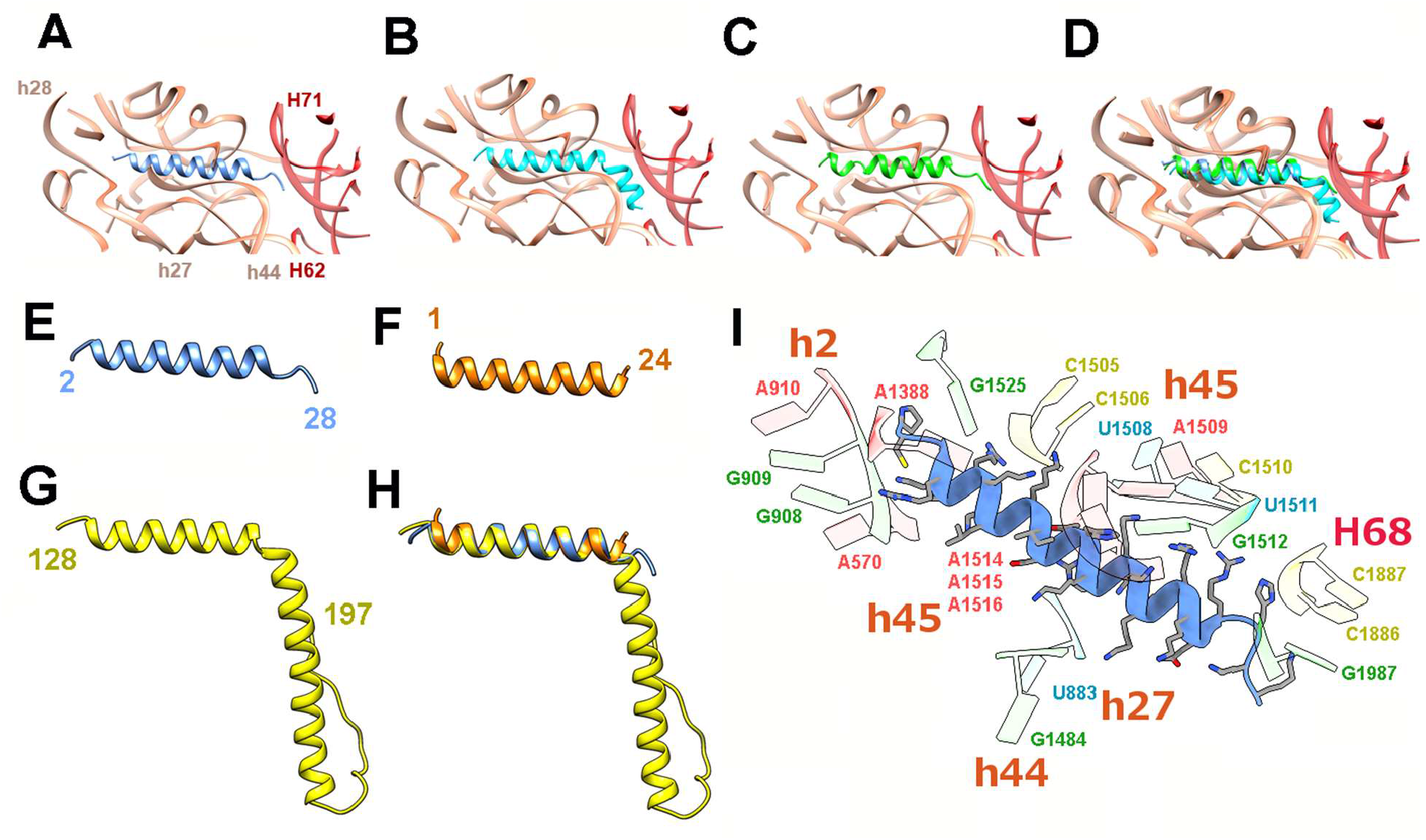
The bS22 protein structure in *Bbu* and other organisms. **(A)** *Bbu* bS22 (blue) in the *Bbu* ribosome binding pocket; **(B)** *Msm* bS22 (cyan) in the *Msm* ribosome binding pocket; **(C)** *Fjo* bS22 (green) in the *Fjo* ribosome binding pocket; **(D)** Overlay of all three bS22 proteins and their respective binding pockets showing the high structural similarity in the three binding pockets with 16S RNA shown in orange and 23S RNA shown in red; (E) *Bbu* bS22 protein; (F) *Homo sapiens* (*Hsa*) eL41 protein; (G) *Hsa* mitochondrial mS38 protein; (H) overlay of *Bbu* bS22, *Hsa* eL41, *Hsa* mS38 proteins; (I) Molecular interactions of *Bbu* bS22 protein with the ribosomal RNA components of the binding pocket, interacting 16S RNA helix numbers in orange, interacting 23S RNA helix numbers in red.

The *Bbu* ribosome resembles the *Fjo* ribosome more than the *Msm* ribosome in another aspect – the presence of the bS21 protein. The bS21 protein in the *Fjo* ribosome, along with the proteins bS6 and bS18, are proposed to help form a binding site for sequestering the Anti-Shine Dalgarno (ASD) sequence for efficient translation of mRNA transcripts that do not contain a Shine-Dalgarno (SD) sequence^20^. This binding pocket is also present in the *Bbu* ribosome but has some differences. The bS21 protein’s C-terminal end in *Bbu* is longer than that in *Fjo* and forms a well-ordered longer helix (Figure S4). The bS6 C-terminal end is also longer in *Bbu* but like the ASD sequence region of *Bbu* 16S RNA is not resolved in the cryo-EM density map. However, the bS6 C-terminal end does show a propensity to form a short helix like the *Fjo* bS6 in the predicted Alphafold model of full-length bS6 (Figure S4D). Only some of the specific residues implicated in stabilizing the ASD sequestration in *Fjo*^20^ are maintained in *Bbu* (*Fjo* bs18 Gln58, Phe50, Leu62, Leu66 are analogous to *Bbu* bs18 Gln59, Phe51, Leu63 and Ile67). Other such residues are altered, e.g., Tyr54 in *Fjo* bS21 is replaced by Lys53 in *Bbu* bs21. Even with these differences, it is possible that this binding pocket in the *Bbu* ribosome can stabilize the ASD sequence in *Bbu* 16S RNA, perhaps to a lesser extent than *Fjo*, which might especially be useful when translating leaderless mRNA transcripts that occur with a high frequency in *Bbu*.

In the *Bbu* ribosome large subunit, there is a structural feature that has not been previously observed in any bacterial ribosome structure. The uL30 protein in *Bbu* (Figure 1C, 1G) is a structural and functional representation of two separate ribosomal proteins – uL30 and bL37 (Figure 4A – 4B, supplementary movie 3). The bL37 protein, a small helical protein that has only been found in mycobacterial ribosomes so far, occupies a pocket near the peptidyl transferase center (PTC). In *Bbu*, there is no separate bL37 protein, but a N-terminal helical extension of uL30 occupies the same pocket by adopting a bent helical structure (Figure 4A, 4C). Helical N-terminal extensions of uL30 are present in non-*Borrelia* bacterial species (Figure S5) as well as in other *Borrelia* species (Figure S6), indicating that this feature is not unique to *Bbu*.

**Figure 4.**
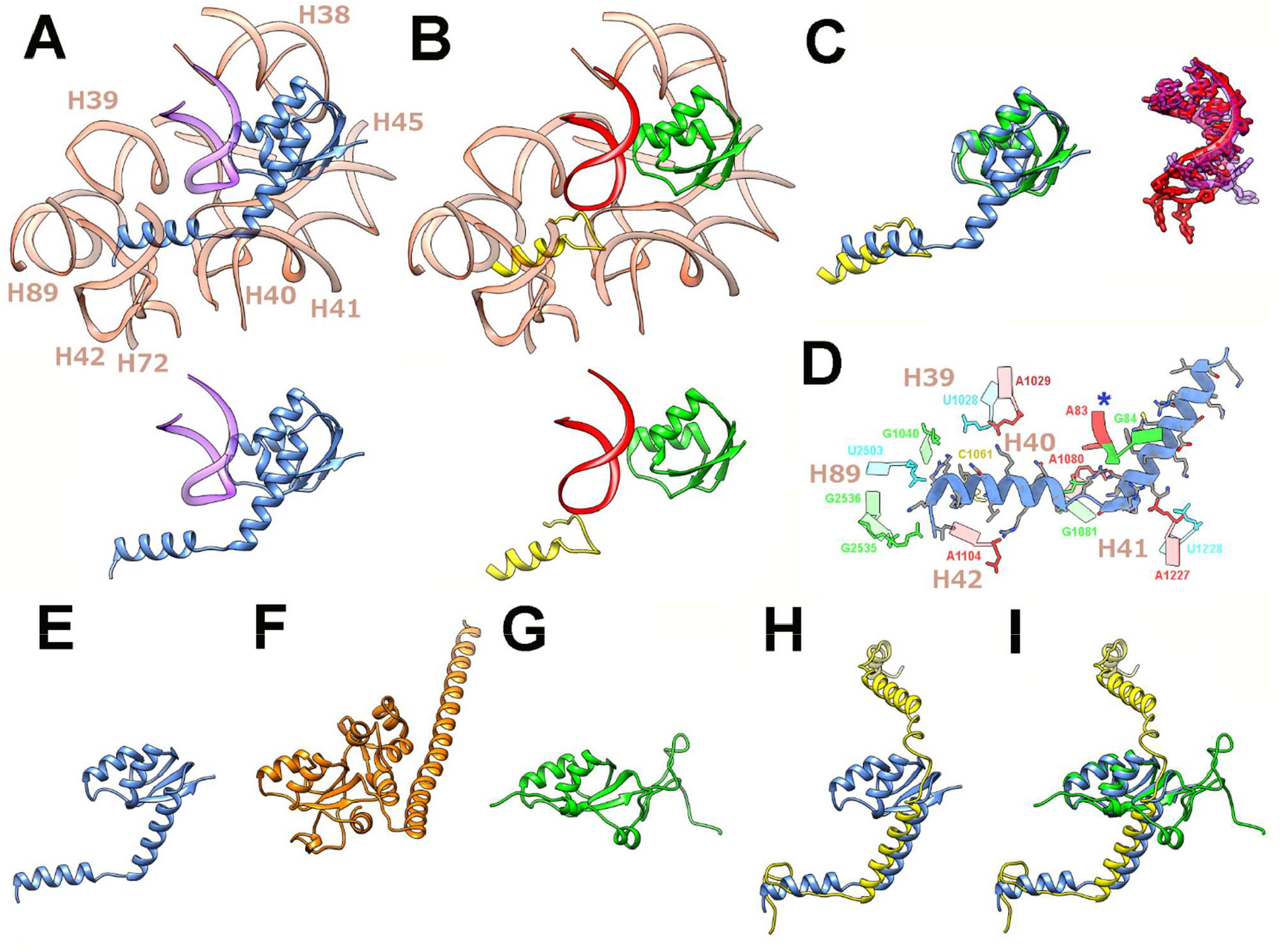
The structure of the longer uL30 protein in *Borreliella* species and its relationship with the bL37 protein in mycobacteria, the *Hsa* uL30 protein, and the *Hsa* mL30 and mL63 proteins. **(A)** The *Bbu* uL30 protein (blue) in its structurally conserved 23S RNA (light orange) and 5S RNA (purple) environment, with the extracted uL30 and 5S RNA shown below for clarity. **(B)** The *Msm* uL30 protein (green) and the *Msm* bL37 protein (yellow) in its structurally conserved 23S RNA (light orange) and 5S RNA (red) environment, with the extracted uL30, bL37, and 5S RNA shown below. **(C)** Overlays of the *Bbu* uL30 protein and the *Msm* uL30 and bL37 proteins (left), and the neighboring *Bbu* (purple) and *Msm* (red) 5S RNA segments (right). **(D)** The ribosomal RNA binding pocket for the uL30 N-terminal extension (residue 2-40), blue asterisk indicates 5S RNA. **(E)** The *Bbu* uL30 protein. **(F)** The *Hsa* uL30 protein. **(G)** The *Hsa* uL30m protein. **(H)** Overlay of the *Bbu* uL30 protein (blue) and the *Hsa* mL63 protein (yellow). **(I)** Overlay of the *Bbu* uL30 protein and the *Hsa* uL30m and *Hsa* mL63 proteins.

The binding pocket accommodating the N-terminal *Bbu* uL30 extension is composed of 23S RNA nucleotide residues from helices H39-H42 and H89, as well as 5S RNA nucleotide residues (Figure 4D). The *Bbu* uL30 protein can also likely be expressed in a shorter form not containing the N-terminal helix before the bend (i. e., the helix corresponding to mycobacterial bL37), which suggests that the occupation of the bL37 pocket could be controlled by regulating the relative expression of these shorter and longer forms of *Bbu* uL30 (Figure S6). The *Bbu* uL30 protein resembles the mammalian cytosolic uL30 protein in having an N-terminal helical extension, except that the uL30 N-terminal helical extension in the mammalian cytosolic ribosome is not bent and is oriented in a different direction (Figure 4F). The *Bbu* uL30 protein also has similarities to mammalian mitochondrial proteins uL30m and mL63, which together occupy the analogous regions in the mammalian mitochondrial ribosomal large subunit (Figure 4G – 4I). These observations suggest the possibility that an N-terminal extension of ul30 might have evolutionarily bifurcated into two proteins in mycobacteria and mammalian mitochondria. Such bifurcation and subsequent expansion of one or both resulting ribosomal proteins may provide a mechanism by which ribosomal protein expansions might have coevolved in mammalian mitochondrial ribosomes in conjunction with ribosomal RNA reductions.

The *Bbu* ribosomal large subunit also has a clear density corresponding to the recently identified bL38 protein in the *Fjo* ribosome structure (Figure 1C, 1F). A ribosomal protein at this location was not expected to be found in *Bbu* as there is no annotated bL38 protein in the *Bbu* genome. In fact, we have not yet been able to identify a protein sequence corresponding to this density in the *Bbu* genome through translated nucleotide searches or mass spectrometric identification of protein components of the *Bbu* ribosome. The density is strong enough to create a reliable backbone model and has some clear sidechain densities that will help distinguish between candidate sequences but our attempts to use *de novo* methods to identify a sequence for this protein through the cryo-EM density alone have not been successful.

Three-dimensional (3D) classification of the cryo-EM particle images did not yield clearly different structural populations, which suggests that our sample of the hibernating *Bbu* 70S ribosome was mostly homogeneous (Figure S7). Some variability was observed in the presence or absence of parts of the bS2 protein density, but no individual class could be identified with a complete bS2 density present. A small number of particles could be separated into a class mostly corresponding to the 50S subunit with a miniscule proportion of 70S particles within it that could not be eliminated entirely. This class of 12,449 particles was used to reconstruct a *Bbu* 50S subunit structure at a resolution of 3.4 Å (Figure S8A). The protein densities corresponding to the full-length uL30 and bL38 proteins were found unaltered in the 50S subunit structure (Figure S8 B-E), confirming that these proteins are stable components of the 50S subunit.

A surprising finding in this 50S subunit structure was that the long and usually well-resolved 23S RNA helix H68 was not even partially ordered (Figure S9). This disorder, also present for the usually flexible H69, seems to be greater than the alternate conformations identified for H68 in the *Staphylococcus aureus* (*Sau*) 50S subunit and 70S ribosome at physiological temperature (Figure S10, supplementary movie 4) that were suggested to be involved in the ribosomal translocation mechanism^40^. A lowering of the density threshold shows some ordering of H68 but only with simultaneous appearance of the 30S subunit density, suggesting that this ordering may be attributable to the small number of 70S particles in this class. One explanation for the H68 disorder is that the 70S sucrose gradient fraction has some not-fully-assembled 50S population. Another explanation is that this 50S population is formed through splitting of a small 70S sub-population. H68 having a larger conformational variability than previously found would be incompatible with proper 70S structure maintenance. The nearby inter-subunit bridge B7a near the L1 stalk region may be one of the final ones to dissociate during ribosomal subunit splitting^41^ and larger scale H68 motions could affect this bridge. With such larger motions, the H68 helix could therefore play a role in completing the dissociation of the two ribosomal subunits. If so, the lack of any special features in H68 in *Bbu* as compared to other bacterial species suggests that this postulated ribosome splitting assistive role for H68 might be common to other ribosomes as well.

This *Bbu* 70S ribosome structure captures details of antibiotic binding pockets that allow construction of detailed models of antibiotics bound to it. We have modeled the structure of three antibiotics, two that are in clinical use already (doxycycline and erythromycin), and one that has recently been shown to be of great promise in treating Lyme infections (hygromycin A^22^) using *de novo* docking (Figure S11) and structural analogy to previous antibiotic-bound ribosome structures (Table S5, Figure S12). The *de novo* docking, done in the vicinity of the expected binding sites, found binding poses analogous to the previous experimental structures with the Autodock Vina^42^ estimated binding free energies of -5.4 kcal/mol for doxycycline, -6.2 kcal/mol for erythromycin, and -7.3 kcal/mol for hygromycin A. A comparison between the hygromycin A binding site for *Bbu* and the same site in the *Tth* ribosome, with^43^ and without^25^ hygromycin A bound, reveals that some 23S RNA residues in *Bbu* are intrinsically positioned in conformations forming a more open binding pocket conducive to hygromycin A binding (Figure S13). Recent studies have shown that antibiotics may perform their function by introducing protein synthesis malfunctions in a context-specific manner^44,45^. In their empty ribosome binding modes, these antibiotics sterically overlap with components that bind to the translating ribosomes suggesting that they may adjust their ribosome binding modes to the presence of these other components. Structurally characterizing *Bbu* translation factors and *Bbu* tRNAs bound to its ribosome in various steps of protein translation, in the presence and absence of antibiotics, can help clarify their context-specific mechanism of action.

In summary, this study identifies unanticipated alterations in the *Bbu* ribosome, such as the presence of bS22, bL38, and an N-terminal extended uL30 also assuming the role of the bL37 protein. The extended uL30 being present in ancient bacterial species (Figure S5) as well as having eukaryotic cytosolic and mitochondrial ribosome analogies (Figure 4D - 4H) suggests this to be analogous to an earlier version of uL30 in evolution that has subsequently shortened or split or expanded, which suggests an underlying mechanism for the increase in ribosomal protein numbers and masses in eukaryotic mitochondrial and cytosolic ribosomes. Our hibernating *Bbu* 70S ribosome structure thus provides the groundwork for better understanding ribosome dormancy, ribosome evolution, antibiotic mechanisms of action, and for development of new structure-based antibiotic therapeutics for Lyme disease.

**Supplementary material.** Text description sections 1.1-1.2, Tables S1-S4, and Figures S1-S13 are provided in a supplementary document. Two svg format files are provided depicting the secondary structures of the 23S RNA, 5S RNA, and 16S RNA derived from our 70S *Bbu* ribosome structure. Four supplementary movies are included that show features of the 70S ribosome structure, relative motion of HPF and E-tRNA, analogy of *Bbu* uL30 protein to other ribosomal proteins, and disorder or alternate conformations for the H68 helix in 23S RNA for *Bbu* and *Sau*.

## Methods

### Ribosome purification

The *Bbu* strain B31-A3^46^ was inoculated in 2L BSK II complete medium and grown to the late logarithmic phase. Cells were harvested and lysed by french press for two pressings at 16,000 PSI and the lysate centrifuged twice at 10,000 rpm for 30 minutes. Formation of a pellet of spirochete cells, expected to occur after spinning at 8,000 rpm, was observed after the first spin. The second spin yielded a much smaller pellet. The supernatant after the second spin was examined by dark microscopy at 400X magnification for intact spirochetes. After observing cell debris from approximately three dead spirochetes in the lysate, a final spin of 17,000 rpm for 45 minutes was done to which ensured pelleting of all the spirochete cell debris. The supernatant was collected in a Beckman PC ultracentrifuge tube and centrifuged for 2 hours and 15 minutes at 42,800 rpm in a Beckman rotor Type 70Ti. Pellets were soaked in 2 mL of HMA-10 buffer (20 mM HEPES K pH 7.5, 600 mM NH4Cl, 10 mM MgCl2, 5 mM β-mercaptoethanol) and kept in an ice bath overnight. 16 mL of HMA-10 buffer was added and the preparation was put on a shaking rocker for one hour at 4 °C with 3 units/mL Rnase-free Dnase (Ambion) added. The contents were transferred to Beckman PA tubes and centrifuged at 13,500 rpm at 4 °C for 15 minutes in Beckman rotor JA 30.5Ti. The supernatant was collected in Beckman ultracentrifuge PC tubes and centrifuged for 2 hours and 15 minutes at 4°C at 42,500 rpm in a Beckman rotor Type 70Ti. Pellets were resuspended in 200 μL HMA-10 buffer and kept in an ice bath in a cold room at 4°C overnight. The sample was then centrifuged for 6 minutes at 13,000 rpm. The supernatant containing the ribosomes was collected and quantified by measuring optical density at 260nm. This crude ribosome preparation was then layered on top of sucrose density gradients (10%-40%), prepared in 11 mL tubes containing HMA-10 buffer for 17 hours at 18,000 rpm in a Beckman rotor SW 41Ti. Ribosome fractions containing primarily 70S monosomes were collected after fractionating the sucrose gradient in a 260nm Teledyne ISCO gradient analyzer. These pooled 70S fractions were pelleted by ultracentrifugation at 42,800 rpm for 6 hours in a Beckman rotor Type 70Ti, suspended in HMA-10 buffer, and quantified by measuring absorbance at 260 nm.

### Grid preparation and imaging for cryo-EM

A home-made thin carbon film was coated as a continuous layer (about 50 Å thick) onto Quantifoil 300-mesh 1.2/1.3 grids. These grids were then glow-discharged for 30 seconds on a plasma sterilizer to hydrophilize the carbon film. The purified ribosome sample (4 μL) was transferred onto each grid after mounting it on a Thermofisher Vitrobot Mark IV system and the grid was maintained for 15 seconds at 4 °C and 100% humidity. Each grid was then blotted for 5 seconds with a force offset of +2 and then plunged into liquid ethane. Movies were collected in counting mode using Leginon software^47^ on a Thermofisher Titan Krios G3 electron microscope operating at 300 kV with a Gatan BioQuantum imaging energy filter and a Gatan K2 direct electron detection (DED) camera. The cryo-EM data collection details are shown in Table 2. A total of 4,661 movies, each having 50 frames, each frame collected every 0.2 seconds, were obtained at a physical pixel size of 1.0961 Å. An electron dose rate of about 8.1 electrons/pixel/second and an exposure time of 10 seconds yielded a total dose of 67.5 electrons/Å^2^.

**Table 2.**
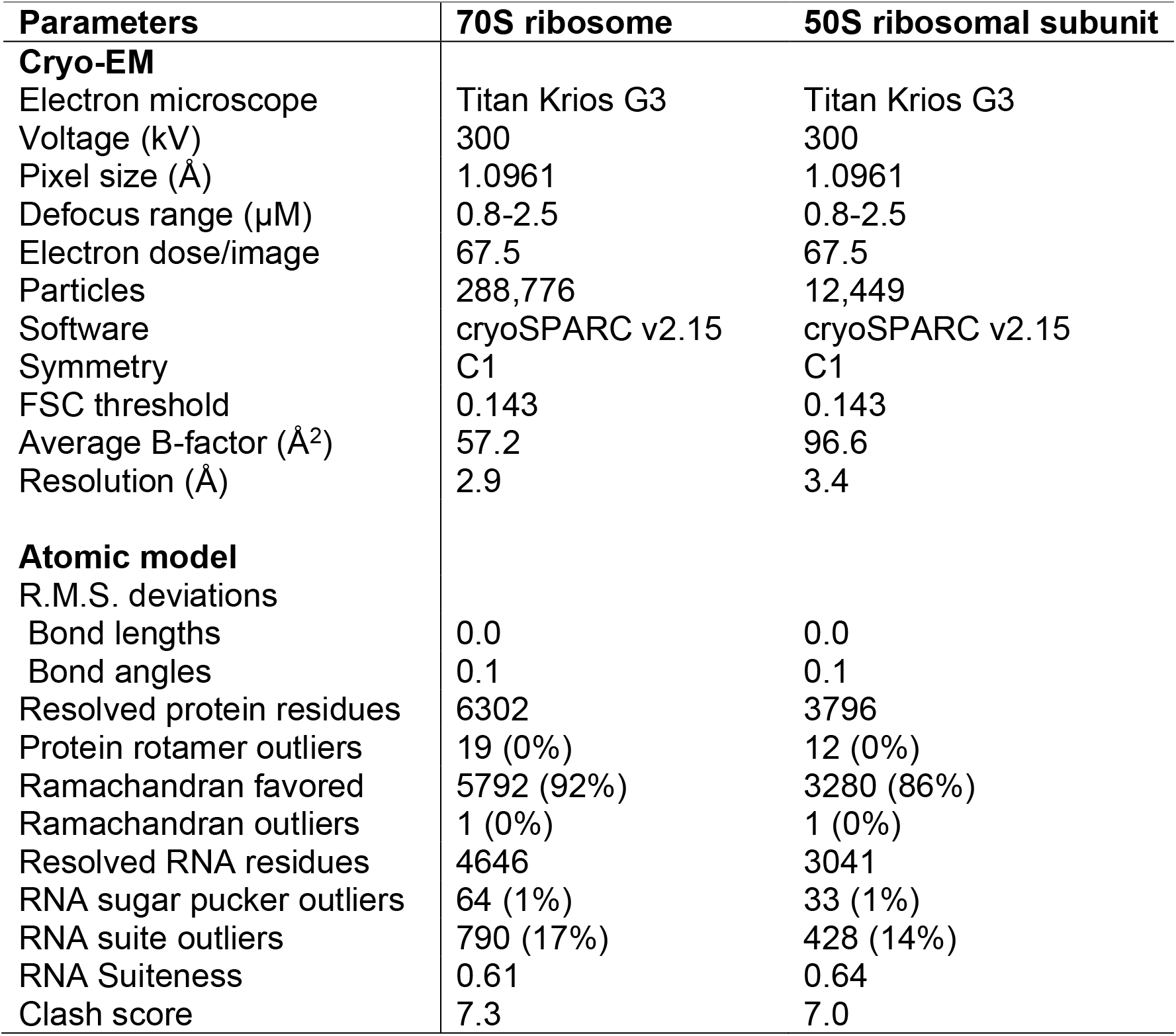
Cryo-EM data collection and model refinement details.

| Parameters | 70S ribosome | 50S ribosomal subunit |
| --- | --- | --- |
| **Cryo-EM** |  |  |
| Electron microscope | Titan Krios G3 | Titan Krios G3 |
| Voltage (kV) | 300 | 300 |
| Pixel size (Å) | 1.0961 | 1.0961 |
| Defocus range (μM) | 0.8-2.5 | 0.8-2.5 |
| Electron dose/image | 67.5 | 67.5 |
| Particles | 288,776 | 12,449 |
| Software | cryoSPARC v2.15 | cryoSPARC v2.15 |
| Symmetry | C1 | C1 |
| FSC threshold | 0.143 | 0.143 |
| Average B-factor (Å<sup>2</sup>) | 57.2 | 96.6 |
| Resolution (Å) | 2.9 | 3.4 |
| **Atomic model** |  |  |
| R.M.S. deviations |  |  |
| Bond lengths | 0.0 | 0.0 |
| Bond angles | 0.1 | 0.1 |
| Resolved protein residues | 6302 | 3796 |
| Protein rotamer outliers | 19 (0%) | 12 (0%) |
| Ramachandran favored | 5792 (92%) | 3280 (86%) |
| Ramachandran outliers | 1 (0%) | 1 (0%) |
| Resolved RNA residues | 4646 | 3041 |
| RNA sugar pucker outliers | 64 (1%) | 33 (1%) |
| RNA suite outliers | 790 (17%) | 428 (14%) |
| RNA Suiteness | 0.61 | 0.64 |
| Clash score | 7.3 | 7.0 |

### Data processing for cryo-EM

Movies were processed using patch motion and patch contrast transfer function (CTF) correction implemented in cryoSPARC v2.15 ^48^. The micrographs were curated to remove those with CTF fits worse than 5 Å and those with relative ice thickness greater than 1.05. Template-based auto-picking of particles was done using the template picker module in cryoSPARC with the input 2D templates (100 equally spaced views) obtained from an earlier 3.9 Å resolution reconstruction volume of the hibernating *Bbu* ribosome generated from data collected on the local JEOL3200FSC 300 KV electron microscope using our automation protocol for SerialEM^49^. The automated particle picks were filtered for particles with normalized correlation coefficient (NCC) > 0.2 and a signal amplitude between 3175 and 5454, which selected a total of 541,319 particles and excluded 98,553 particles. The selected particles were extracted from the corrected micrographs at a box size of 380 pixels and then further filtered using multiple iterations of 2D classifications, each followed by selection of good particle classes prior to using them as input for the next 2D classification, yielding a final selection of 288,776 particles used for further processing. These particles were used for generating a 3D reconstruction using *ab initio* reconstruction, homogeneous refinement, non-uniform refinement^50^, and finally local non-uniform refinement with the mask from the previous refinement, whose final gold-standard 3D refinement resolution was 2.9 Å. Further classification was attempted using 3D variability analysis, heterogeneous refinement, and *ab initio* reconstruction with multiple classes, but the resulting classes showed only small differences from each other, such as presence or absence of fragmented partial density of uS2 protein, suggesting that the particles mostly represented a homogeneous population. The only exception was a small class of 12,449 particles consisting of mostly 50S subunit particles with a slight contamination of 70S particles, whose final gold-standard resolution was 3.4 Å.

### Model building and refinement

The initial models for the 23S, 16S and 5S ribosomal RNAs were built by manually aligning the *Bbu* sequences to the corresponding sequences of the mycobacterial^18^ ribosomal RNAs and using RNA homology modeling program Moderna ^51^ to generate initial homology models using a known RNA structure (PDB ID: 5O61) ^18^. The RNA secondary structures for the *Bbu* 23S, 16S and 5S ribosomal RNAs were obtained from their modeled 3D structures using the RNA PDBEE server^52,53^ and, with the aid of the R2DT server^54^, were manually converted to the scalable vector graphic (svg) format depictions that are provided in the supplementary material. Initial atomic models for most *Bbu* ribosomal proteins and bbHPF were built using Alphafold2^55^. The only exception is the non-annotated bL38 protein for which no corresponding genomic sequence has yet been identified. It has therefore been built manually into the density as a backbone only model. Manual rebuilding of incorrectly modeled regions was performed using UCSF Chimera v1.16 ^56^ followed by restrained local real-space refinement in Phenix v1.18 ^57^. UCSF Chimera v1.16 ^56^ or UCSF ChimeraX version 1.4^58,59^ were used to make the molecular figures, Inkscape was used to edit the RNA secondary structure depictions, and Gimp v2.10 was used to make composite figures.

## Supporting information

23srna_5srna_secondary_structure_svg

16srna_secondary_structure_svg

## Data availability

Cryo-EM volumes and atomic models have been deposited at the EMDB (accession codes: EMD-29298, EMD-29304) and PDB (accession codes: 8FMW, 8FN2), for the 70S ribosome and the 50S subunit, respectively.

## Code availability

The programs used for cryo-EM image processing, structure determination, refinement, and analysis are all publicly available.

## Acknowledgements

We acknowledge the use of the Wadsworth Center cryo-EM facility and the help and training provided by Chyongere Hsieh and Michael Marko for this use. We thank Joseph Wade for discussions that helped identify the bS22 protein sequence. The cryo-EM data was collected at the Simons Electron Microscopy Center and National Resource for Automated Molecular Microscopy located at the New York Structural Biology Center, supported by grants from the Simons Foundation (349247), NYSTAR, and the NIH National Institute of General Medical Sciences (GM103310). This work was supported by the NIH NIGMS grant (GM061576) to R.K.A. R.K.A. also acknowledges support to his lab through NIH R01 grants AI132422, GM139277 and AI155473.

## Contributions

M.R.S., Y-P.L., R.K.A., and N.K.B. designed the project. A.L.M. grew the *Bbu* culture, lysed the cells, and ensured spirochete cellular disintegration using light microscopy. P.K. purified the *Bbu* ribosomes. S.M. analyzed the proteins in the ribosome preparation. R.K.K. prepared the cryo-EM grids. R.K.K. and N.K.B. did the electron microscopy at Wadsworth Center. N.K.B. processed the cryo-EM data to generate final 3D volumes. M.R.S., E.K.A., and N.K.B. built protein models. S.R.M. and N.K.B. built ribosomal RNA and tRNA models. N.K.B. generated and refined the final combined atomic model built into the 3D volumes and performed the initial structural analysis. M.R.S., S.R.M., R.K.K., S.M., R.K.A., and N.K.B. contributed to further structural analysis and all authors participated in writing or editing the manuscript.

