## supplementary_material_document for "The structure of a hibernating ribosome in a Lyme disease pathogen"

a. Present address – University of California at Berkeley, Berkeley, CA

b. Present address – Robert P. Apkarian Integrated Electron Microscopy Core, Emory University, Atlanta, GA

c. Present address – National Center for Biological Sciences, Bangalore, India

**1.1 Finding the bS22 ribosomal protein in *Borrelia (Borreliella) burgdorferi* (*Bbu*).**

A small helical density in our *Bbu* 70S ribosome cryo-EM map corresponded to the bS22 protein seen in mycobacterial[1-4] and Bacteroidetes[5] ribosomal small subunit structures, indicating that the bS22 protein is also present in *Bbu*. Since this protein is not annotated in the *Bbu* genome, the mycobacterial bS22 sequence from *Mycobacterium smegmatis* (*Msm*): MGSVIKKRRKRMSKKKHRKLLRRTRVQRRKLGK; was used as a query sequence for a tblastn[6] search of the *Bbu* genome with a word size of 2 and no low complexity filter. This yielded hits for a smaller fragment of the query sequence but no plausible *Bbu* bS22 protein with a size near 30 amino acid (aa) residues upon translation of the genomic hits in 6 frames. The same search was then attempted with the Bacteroidetes bS22 sequence from *Flavobacterium johnsoniae* (*Fjo*): MPSGKKRKRHKVATHKRKKRARANRHKKKK. This search yielded one hit, which when translated in the 6 frames, yielded the plausible bS22 sequence: VPCGRKRKLKKISTHKRKKKRRKNRHKKKNK-, with an appropriately placed stop codon "-". The first valine residue is likely translated as a methionine residue since substitution for Met (AUG) instead of Val (GUG) can occur due to the anticodon for fMet-tRNA (CAU) pairing well enough with GUG on the mRNA through wobble pairing between G and U. Such usage of GUG as a start codon in bacteria is known to occur at an average of 12%[7,8]. This sequence fits well into the bS22 sidechain cryo-EM densities, allowing it to be identified as the correct *Bbu* bS22 sequence.

A ClustalW 2.1[9] pair-wise sequence alignment between *Bbu* and *Fjo* bS22 sequences shows a 65% sequence identity, which explains why the *Fjo* query sequence tblastn search was successful:

```
Bbu            MPCGRKRKLKKISTHKRKKKRRKNRHKKKNK
Fjo            MPSGKKRKRHKVATHKRKKRARANRHKKKK-
               **.*:*** :*::******: * ******:
```

A ClustalW 2.1[9] multiple sequence alignment between *Bbu*, *Fjo*, and *Msm* bS22 sequences shows that identity of residues between the three sequences drops to 19%, which explains why the *Msm* query sequence tblastn search was unsuccessful:

```
Bbu            MPCGRKRKLKKISTHKRKKKRRKNRHKKKNK--
Fjo            MPSGKKRKRHKVATHKRKKRARANRHKKKK---
Msm            MGSVIKKRRKRMSKKKHRKLLRRTRVQRRKLGK
               *  .  *.. :.::.:*..*  * .* :..:
```

The genomic sequence identified for the *Bbu* B31 bS22 protein (Genbank ID: AE000783.1, nucleotides 867605-867697) is:

```
GTGCCTTGCGGAAGAAAAAGAAAATTAAAAAAAATTTCTACCCATAAAAGGAAGAAAAAAAGAA
GAAAAAATAGACATAAGAAAAAAAAATAAG
```

### 1.2 Alignments of proteins that sequester Anti-Shine Dalgarno (ASD) sequence.

#### bS6, Identity: 23/141 (16.3%), Similarity: 50/141 (35.5%)

```
    1        10          20        30        40        50        60
Bbu MIKRYEACFLFKS--EEIEYKGSLEEVKKSLEFFGATDVVSNFIGERALEYPIKKQARGRYEIIEFS
Fjo -MNHYETVFILNPVLSEVQVKETVTKFEEFLTSRGAEMVSKEDWGLKKMAYEIQNKKSGFYHLFEFK
     1        10        20        30        40        50        60
     :..**..*:...  .*::.*.::.:.::.*...**..*.....*.:.:.*.* :.:..*.*.:.**.

    66  70        80        90        100       110       120       130
Bbu MEGNNLKEFESRLKLIRNLLRYMILVKIVRKINTKKIKRRNFREFRDNIDKDSLKGASKVETPTGPE
Fjo VAGEVLIAFETEFRRDERVMRFLTVSLDKHAISWAERRRAKLKSTKA--------------------
    67 70        80        90        100       110
    :.*..*..**:..:....::*::.:......*:..:.:*...:..:.

    133   139
Bbu STDIQEK
Fjo -------
```

#### bS18, Identity: 42/102 (41.2%), Similarity: 64/102 (62.7%)

```
    1        10         20        30         40         50        60
Bbu MYKDRDTNQRDSRFENQQD-GFKKNSNFRFFK-RKSCKFCDSG-KHPDYKDFDFLKKFITEQGKILP
Fjo ----MSTIEQSAKGKKDGDIRYLTPLNIETNKTKKYCRFKKSGIKYIDYKDADFLLKFVNEQGKILP
        1        10        20        30        40        50        60
        ..*.::.::.:...* .:....*....* :*.*:*..** *:.****.***.**:.*******

    65   70        80        90    96
Bbu KRITGTSAKHQRRLALEIKRARYMALLPFVKK---
Fjo RRLTGTSLKYQRKVSVAVKRARHLALMPYVADLLK
    64    70        80        90       98
    :*:****.*:**::::.:****::**:*:*..
```

#### bS21, Identity: 19/70 (27.1%), Similarity: 36/70 (51.4%)

```
    1        10        20        30        40         50        60      68
Bbu MVTVTVDKNENLEKALKRFKRMIEKEAIIREWKRREYYEKPSTI-RVKKEKAFKRKQAKKVRKLKQKTNR
Fjo MLIIPIKDGENIDRALKRYKRKFDKTGTVRQLRARTAFIKPSVVKRAQIQKAAYIQTLKDSLES------
    1        10        20        30        40        50        60  64
    *:.:.:...**:::****:**..:*...:*:.:.*..:.***.: *.:.:**...:..*...:.
```

#### 16S RNA 3’-end

```
Bbu UCGUAACAAGGUAGCCGUACUGGAAAGUGCGGCUGGAUCACCUCCUUU--
Fjo UCGUAACAAGGUAGCCGUACCGGAAGGUGCGGCUGGAACACCUCCUUUCU
    ******************** **** *********** **********
```

Core ASD sequence

**Table S1. The resolved 58 components of the *Bbu* 70S ribosome and 50S subunit.**

| Name | 8FMW Chain ID | 8FN2 (50S) Chain ID | Size (residues) | Modeled (residues) | Accession ID* | Comments |
|---|---|---|---|---|---|---|
| 23S RNA | AA | A | 2933 | 4-2932 | AE000783.1 | Nucleotides: 438267-435336 |
| 5S RNA | AB | B | 112 | 1-112 | AE000783.1 | Nucleotides: 435312-435201 |
| uL1 | AC | C | 226 | 6-226 | O51353 | Backbone model |
| uL2 | AD | D | 277 | 1-277 | P94270 | |
| uL3 | AE | E | 206 | 1-206 | P94267 | |
| uL4 | AF | F | 209 | 1-209 | P94268 | |
| uL5 | AG | G | 182 | 1-182 | O51443 | |
| uL6 | AH | H | 180 | 1-180 | O51446 | |
| bL9 | AI | I | 173 | 1-148 | O51139 | Partial backbone model (41-148) |
| uL10 | AJ | J | 162 | 1-162 | O51352 | |
| uL11 | AK | K | 143 | 5-143 | O51354 | |
| uL13 | AL | L | 146 | 2-146 | O51314 | |
| uL14 | AM | M | 122 | 1-122 | O51441 | |
| uL15 | AN | N | 145 | 1-145 | O51450 | |
| uL16 | AO | O | 138 | 1-138 | O51438 | |
| bL17 | AP | P | 123 | 1-121 | O51456 | |
| uL18 | AQ | Q | 119 | 1-119 | O51447 | |
| bL19 | AR | R | 121 | 1-117 | O51642 | |
| bL20 | AS | S | 115 | 2-115 | O51206 | |
| bL21 | AT | T | 103 | 1-103 | O51719 | |
| uL22 | AU | U | 120 | 2-116 | P94272 | |
| uL23 | AV | V | 98 | 1-98 | P94269 | |
| uL24 | AW | W | 101 | 1-101 | O51442 | |
| bL25 | AX | X | 182 | 2-182 | O51727 | |
| bL27 | AY | Y | 81 | 8-81 | O51721 | |
| bL28 | AZ | Z | 92 | 2-92 | O51325 | |
| uL29 | Aa | a | 65 | 1-65 | O51439 | |
| uL30 | Ab | b | 101 | 2-101 | O51449 | 2 possible translation products |
| bL31 | Ac | c | 81 | 1-81 | O51247 | |
| bL32 | Ad | d | 60 | 2-60 | O51646 | Zn ion |
| bL33 | Ae | e | 59 | 9-59 | O51357 | |
| bL34 | Af | f | 51 | 1-50 | P29220 | |
| bL35 | Ag | g | 66 | 1-66 | O51207 | |
| bL36 | Ah | h | 37 | 1-37 | O51452 | Zn ion |
| bL38 | Ai | i | Not known | 1-46 | Unidentified | Backbone model |
| E-tRNA | X | - | | | - | Mixture of tRNA populations |
| 16S RNA | A | - | 1538 | 1-1529 | AE000783.1 | Nucleotides: 444581-446118 |
| uS3 | C | - | 293 | 6-226 | P94273 | 67 C-terminal residues not seen |
| uS4 | D | - | 208 | 1-208 | O51560 | |
| uS5 | E | - | 165 | 8-165 | O51448 | |
| bS6 | F | - | 139 | 1-97 | O51142 | 42 C-terminal residues not seen |
| uS7 | G | - | 157 | 1-157 | O51347 | |
| uS8 | H | - | 132 | 1-132 | O51445 | |
| uS9 | I | - | 136 | 6-136 | O51313 | |
| uS10 | J | - | 103 | 2-103 | P94266 | |
| uS11 | K | - | 130 | 14-130 | O51454 | 13 N-terminal residues not seen |
| uS12 | L | - | 124 | 1-124 | O51348 | |
| uS13 | M | - | 125 | 1-114 | O51453 | 11 C-terminal residues not seen |
| uS14 | N | - | 61 | 2-61 | O51444 | Zn ion |

| | | | | | | |
|---|---|---|---|---|---|---|
| uS15 | O | - | 88 | 1-88 | O51744 | |
| bS16 | P | - | 86 | 1-83 | O51638 | |
| uS17 | Q | - | 84 | 3-84 | O51440 | |
| bS18 | R | - | 96 | 34-96 | O51140 | Zn ion, first 33 residues not seen |
| uS19 | S | - | 92 | 2-85 | P94271 | |
| bS20 | T | - | 85 | 1-85 | P49394 | |
| bS21 | U | - | 69 | 1-69 | O51271 | |
| bS22 | V | - | 31 | 2-28 | AE000783.1 | Nucleotides: 867605-867697 |
| bbHPF | W | - | 97 | 1-97 | O51405 | non-ribosomal protein |

*Accession IDs are Genbank for RNA and Uniprot for proteins (except for the unannotated 30S ribosomal protein bS22). 30S ribosomal proteins uS1 and uS2 are present but uS1 is not resolved and uS2 is only partially resolved at very low threshold and therefore is not modeled.

**Table S2. Details of bacterial HPF structures with resolutions better than 3.5 Å that were used for sequence and structural comparisons.**

| PDB ID | Res. (Å) | Method | Year | Species (R, HPF) | Ref. | Component notes |
|---|---|---|---|---|---|---|
| 4HEI | 1.6 | X-ray | 2013 | *-, Vch* | [10] | HPF alone, no ribosome |
| 4Y4O | 2.3 | X-ray | 2015 | *Tth, Eco* | [11] | YfiA |
| 6S0X | 2.4 | Cryo-EM | 2019 | *Sau, Sau* | [12] | HPF, erythromycin |
| 7RQA | 2.4 | X-ray | 2022 | *Tth, Eco* | [13] | YfiA, tRNA analogs |
| 7RQE | 2.4 | X-ray | 2022 | *Tth, Eco* | [13] | YfiA, tRNA analogs, chloramphenicol |
| 3TQM | 2.5 | X-ray | 2015 | *-, Cbu* | [14] | HPF alone, no ribosome |
| 7RQB | 2.5 | X-ray | 2022 | *Tth, Eco* | [13] | YfiA, tRNA analogs |
| 7RQC | 2.5 | X-ray | 2022 | *Tth, Eco* | [13] | YfiA, tRNA analogs |
| 7RQD | 2.5 | X-ray | 2022 | *Tth, Eco* | [13] | YfiA, tRNA analogs, chloramphenicol |
| 2YWQ | 2.6 | X-ray | 2008 | *-, Tth* | [u] | HPF alone, no ribosome |
| 6CFL | 2.6 | X-ray | 2018 | *Tth, Eco* | [15] | YfiA, D-lysyl-CAM |
| 6XHX | 2.6 | X-ray | 2021 | *Tth, Eco* | [16] | YfiA, A2058 unmethylated, erythromycin |
| 4V8I | 2.7 | X-ray | 2011 | *Tth, Eco* | [17] | YfiA |
| 6CFK | 2.7 | X-ray | 2018 | *Tth, Eco* | [15] | YfiA, D-histidyl-CAM |
| 5FDV | 2.8 | X-ray | 2016 | *Tth, Eco* | [18] | YfiA, Pyrrhocoricin |
| 7MD7 | 2.8 | X-ray | 2021 | *Tth, Eco* | [19] | YfiA, chloramphenicol |
| 5FDU | 2.9 | X-ray | 2016 | *Tth, Eco* | [18] | YfiA, Metalnikowin I |
| 5NGM | 2.9 | Cryo-EM | 2017 | *Sau, Sau* | [20] | HPF, 70S from 100S |
| 6Y69 | 2.9 | Cryo-EM | 2020 | *Eco, Eco* | [21] | HPF, TetracenomycinX |
| 7M4Z | 2.9 | Cryo-EM | 2021 | *Aba, Aba* | [22] | HPF, eravacycline |
| 6H4N | 3.0 | Cryo-EM | 2018 | *Eco, Eco* | [23] | HPF, 70S from 100S |
| 4V8H | 3.1 | Cryo-EM | 2012 | *Tth, Eco* | [17] | HPF |
| 6FKR | 3.2 | X-ray | 2018 | *Tth, Eco* | [24] | YfiA, Tur1A |
| 5V8I | 3.3 | X-ray | 2018 | *Tth, Eco* | [u] | YfiA, no uS17 |
| 6GZQ | 3.3 | Cryo-EM | 2018 | *Tth, Tth* | [25] | HPF |
| 5ZEP | 3.4 | Cryo-EM | 2018 | *Msm, Msm* | [4] | HPF |
| 6DZI | 3.5 | Cryo-EM | 2018 | *Msm, Msm* | [26] | HPF (MPY) |

Res. – Resolution, Ref. – Reference number, u - Unpublished structure, Species (R, HPF) refers to the species name abbreviation for ribosome (R) and HPF. Species name abbreviations are as follows: *Vch – Vibrio cholerae*, *Tth – Thermus thermophilus*, *Eco – Escherichia coli*, *Sau – Staphylococcus aureus*, *Cbu – Coxiella burnetii*, *Aba – Acinetabacter baumannii, Msm – Mycobacterium smegmatis*.

**Table S3. Structure-based pair-wise sequence alignment with bbHPF structure for known bacterial HPF structures.**

| ID | Aligned sequence |
|---|---|
| | β1 α1 β2 β3 β4 α2 |
| | bbb bbb aaaaaaaaaaaaa aaa bbbbbbb bbbbbbbb bbbbbbbb aaaaaaaaaaaaaaaaaaaaaaa |
| *Bbu* | ----MEPKIQ-TVNYSLNENEKNFILKKLEKFDTHIKKHIDNLKITIKKEH-----ELFKLDAHIHFN-W-GKIIHIREDGKILLNLIDSAIARLYKTATKEKEKKNNK---- |
| *Sau* | ----IRFEIH-GDNLTITDAIRNYIEEKIGK-LERYFNDVPNAVAHVKVKTYSN--SATKIEVTIPLK---NVTLRAEERNDDLYAGIDLINNKLERQVRKYKTRINRKSRDR |
| *Tth* | ---MNIYKLI-GRNLEITDAIRDYVEKKLAR-LDRYQDGELMAKVVLSLAGSPHVEKKARAEIQVDLP---GGLVRVEEEDADLYAAIDRAVDRLETQVKRFRERRYVGKRHS |
| *Eco1* | -----TMNIT-SKQMEITPAIRQHVADRLAK-LEKWQTHLINPHIILSKEP-----QGFVADATINTP---NGVLVASGKHEDMYTAINELINKLERQLNKLQHKGEARRAA- |
| *Eco2* | ----MQLNIT-GNNVEITEALREFVTAKFAK-LEQYFDRINQVYVVLKVEK-----VTHTSDATLHVN---GGEIHASAEGQDMYAAIDGLIDKLARQLTKHKDKLKQH---- |
| *Cbu* | ----MHIQMT-GQGVDISPALRELTEKKLHR-IQPCRDEISNIHIIFHINK-----LKKIVDANVKLP---GSTINAQAESDDMYKTVDLLMHKLETQLSKYKAK-------- |
| *Vch* | ----MQINIQ-GHHIDLTDSMQDYVHSKFDK-LERFFDHINHVQVILRVEK-----LRQIAEATLHVN---QAEIHAHADDENMYAAIDSLVDKLVRQLNKHKEKL------- |
| *Msm* | ERPHAEVVVK-GRNVEVPDHFRTYVSEKLSR-LERFDKTIYLFDVELDHERNRRQ-RKNCQHVEITARGR-GPVVRGEACADSFYTAFESAVQKLEGRLRRAKDRRKI----- |
| *Aba* | ----MNIEIRTDKNIHNSERLITYVRAELTQEFQRHSERITHFSVHFSDENGDKG-GDKDIHCMIEARPSGLKPVAVHHKAGNIDASIHGAIEKLKRSLEHTFEKKE------ |
| | : .. : . . : : : .. * : |

Sequences aligned using structural alignment tool within UCSF Chimera. *Bbu – Borreliella burgdorferi* (O51405), *Sau - Staphylococcus aureus* (D2Z097), *Tth – Thermus thermophilus* (Q5SIS0), *Eco1 – Escherichia coli* YfiA (P0AD49), *Eco2 – Escherichia coli* HPF (P0AFX0), *Cbu – Coxiella burnetii* (Q83DI6), *Vch – Vibrio cholerae* (H9L4L9), *Msm – Mycobacterium smegmatis* (A0QTK6), *Aba – Acinetabacter baumannii* (V5V8V8). Uniprot IDs for proteins are in parentheses after species name. *Bbu* HPF (bbHPF) numbered secondary structure elements shown on top as α-helix (**a**) or β-strand (**b**). Bacterial HPF sequences are truncated within four residues of the bbHPF sequence. This table is distinct from Table 1 in the main manuscript in showing only residues that are resolved in the 3D structures of the HPFs in the alignment.

### Table S4. Predicted bS22 sequences in other *Borrelia* species.

| Organism | Predicted bS22 protein sequence | Genome ID | Nucleotide Range |
|---|---|---|---|
| **Lyme borreliae** | | | |
| *Borrelia burgdorferi* | MPCGRKRKLKKISTHKRKKKRRKNRHKKKNK | AE000783.1 | 867605-867697 |
| *Borrelia afzelii* | MPCGRKRKLKKISTHKRKKKRRKNRHKKKNK | CP042238.1 | 39086-39178 |
| *Borrelia garinii* | MPCGRKRKLKKISTHKRKKKRRKNRHKKKNK | CP059009.1 | 871181-871273 |
| *Borrelia valaisiana* | MPCGRKRKLKKISTHKRKKKRRKNRHKKKNK | CP009117.1 | 867482-867574 |
| *Borrelia mayonii* | MPCGRKRKLKKISTHKRKKKRRKNRHKKKNK | CP015780.1 | 868560-868652 |
| *Borrelia maritima* | MPCGRKRKLKKISTHKRKKKRRKNRHKKKNK | CP044535.1 | 865321-865413 |
| *Borrelia chilensis* | MPCGRKRKLKKISTHKRKKKRRKNRHKKKNK | CP009910.1 | 864905-864997 |
| *Borrelia miyamotoi* | MPCGRKRKLQKISTHKRKKKRRKNRHKKKNK | CP036914.1 | 39948-40040 |
| **Relapsing fever borreliae** | | | |
| *Borrelia duttonii* | MPCGRKRKLKKISTHKRKKKRRKNRHKKKNK | CP000976.1 | 890138-890230 |
| *Borrelia recurrentis* | MPCGRKRKLKKISTHKRKKKRRKNRHKKKNK | CP000993.1 | 891710-891802 |
| *Borrelia coriaceae* | MPCGRKRKLKKISTHKRKKKRRKNRHKKKNK | CP075076.1 | 876376-876468 |
| *Borrelia crocidurae* | MPCGRKRKLKKISTHKRKKKRRKNRHKKKNK | CP003426.1 | 876767-876859 |
| *Borrelia hermsii* | MPCGRKRKLQKISTHKRKKKRRKNRHKKKNK | CP073148.1 | 879892-879984 |
| *Borrelia parkeri* | MPCGRKRKLQKISTHKRKKKRRKNRHKKKNK | CP073159.1 | 878409-878501 |
| *Borrelia venezuelensis* | MPCGRKRKLQKISTHKRKKKRRKNRHKKKNK | CP073220.1 | 878347-878439 |
| *Borrelia turicatae* | MPCGRKRKLQKISTHKRKKKRRKNRHKKKNK | CP073192.1 | 877560-877652 |
| **Reptile and echidna *Borrelia*** | | | |
| *Candidatus Borrelia tachyglossi* | MPCGRKRKLKKISTHKRKKKRRKNRHKKKNK | CP025785.1 | 894477-894569 |
| | ********* ********************* | | |

All sequences identified through a tblastn[6] search using the *Bbu* bS22 sequence.

**Table S5. Structures of bacterial ribosomes with antibiotics that were used to help generate *Bbu* 70S ribosome antibiotic bound models by structural analogy.**

| PDB ID | Res. (Å) | Method | Year | Species | Ref. | Antibiotic, other notes |
|---|---|---|---|---|---|---|
| 5J5B | 2.8 | X-ray | 2016 | *Eco* | [27] | Tetracycline, |
| 5J7L | 3.0 | X-ray | 2016 | *Eco* | [27] | Tetracycline, at 2 sites, U1052 mutation |
| 4V9A | 3.3 | X-ray | 2013 | *Tth* | [28] | Tetracycline |
| 1HNW | 3.4 | X-ray | 2000 | *Tth* | [29] | Tetracycline |
| 1I97 | 3.5 | X-ray | 2001 | *Tth* | [30] | Tetracycline |
| 6S0Z | 2.3 | Cryo-EM | 2019 | *Sau* | [12] | Erythromycin, ΔR88,ΔA89 uL22 |
| 6S0X | 2.4 | Cryo-EM | 2029 | *Sau* | [12] | Erythromycin, ΔR88,ΔA89 uL22 |
| 6XHX | 2.6 | X-ray | 2021 | *Tth* | [16] | Erythromycin, A2058 unmethylated, YfiA |
| 1YI2 | 2.7 | X-ray | 2005 | *Hma* | [31] | Erythromycin, G2099A |
| 6ND6 | 2.9 | X-ray | 2019 | *Tth* | [32] | Erythromycin, A-, P-, E-tRNAs |
| 7B5K | 2.9 | Cryo-EM | 2021 | *Eco* | [33] | Erythromycin, P-site tRNA nascent chain |
| 7NSO | 2.9 | Cryo-EM | 2021 | *Eco* | [34] | Erythromycin, ErmDL |
| 7Q4K | 3.0 | Cryo-EM | 2022 | *Eco* | [u] | Erythromycin, streptococcal MsrDL |
| 4V7X | 3.0 | X-ray | 2010 | *Tth* | [35] | Erythromycin |
| 4V7U | 3.1 | X-ray | 2010 | *Eco* | [35] | Erythromycin |
| 4WFN | 3.5 | X-ray | 2017 | *Dra* | [36] | Erythromycin, uL22 3-residue insertion |
| 1JZY | 3.5 | X-ray | 2001 | *Dra* | [37] | Erythromycin |
| 7NSP | 3.5 | Cryo-EM | 2021 | *Eco* | [34] | Erythromycin, ErmDL, A-, P-tRNA |
| 5JTE | 3.6 | Cryo-EM | 2016 | *Eco* | [38] | Erythromycin, ErmBL, A-, P-, E-tRNA |
| 5JU8 | 3.6 | Cryo-EM | 2016 | *Eco* | [38] | Erythromycin, ErmBL, P-, E-tRNA |
| 3J7Z | 3.9 | Cryo-EM | 2014 | *Eco* | [39] | Erythromycin, ErmCL |
| 3J5L | 6.6 | Cryo-EM | 2014 | *Eco* | [40] | Erythromycin, ErmBL |
| 5DOY | 2.6 | X-ray | 2015 | *Tth* | [41] | Hygromycin A, A-, P-, E-tRNA |
| 5DM7 | 3.0 | X-ray | 2015 | *Dra* | [41] | Hygromycin A |
| 5DOX | 3.1 | X-ray | 2015 | *Tth* | [42] | Hygromycin A |

Res. – Resolution, Ref. – Reference number, u - Unpublished structure, Species refers to the species name abbreviation for ribosome that are as follows: *Eco – Escherichia coli*, *Tth – Thermus thermophilus*, *Sau – Staphylococcus aureus*, *Hma – Haloarcula marismortui*, *Dra – Deinococcus radiodurans.*

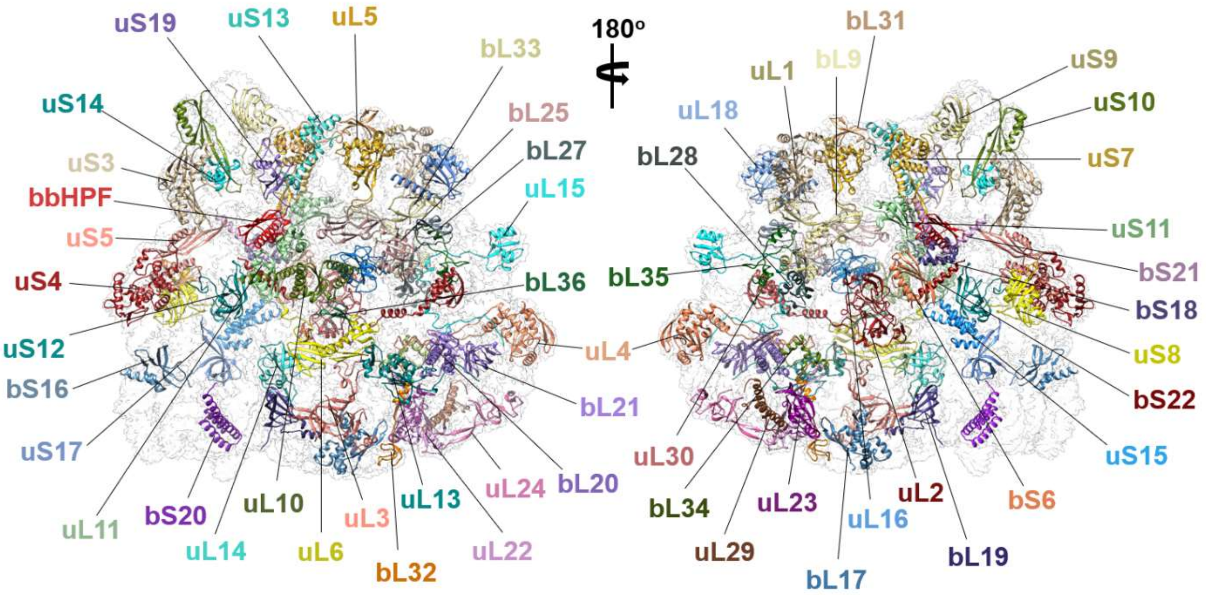


**Figure S1. Protein components of the hibernating *Bbu* 70S ribosome and their labeled models fit in the cryo-EM density.**

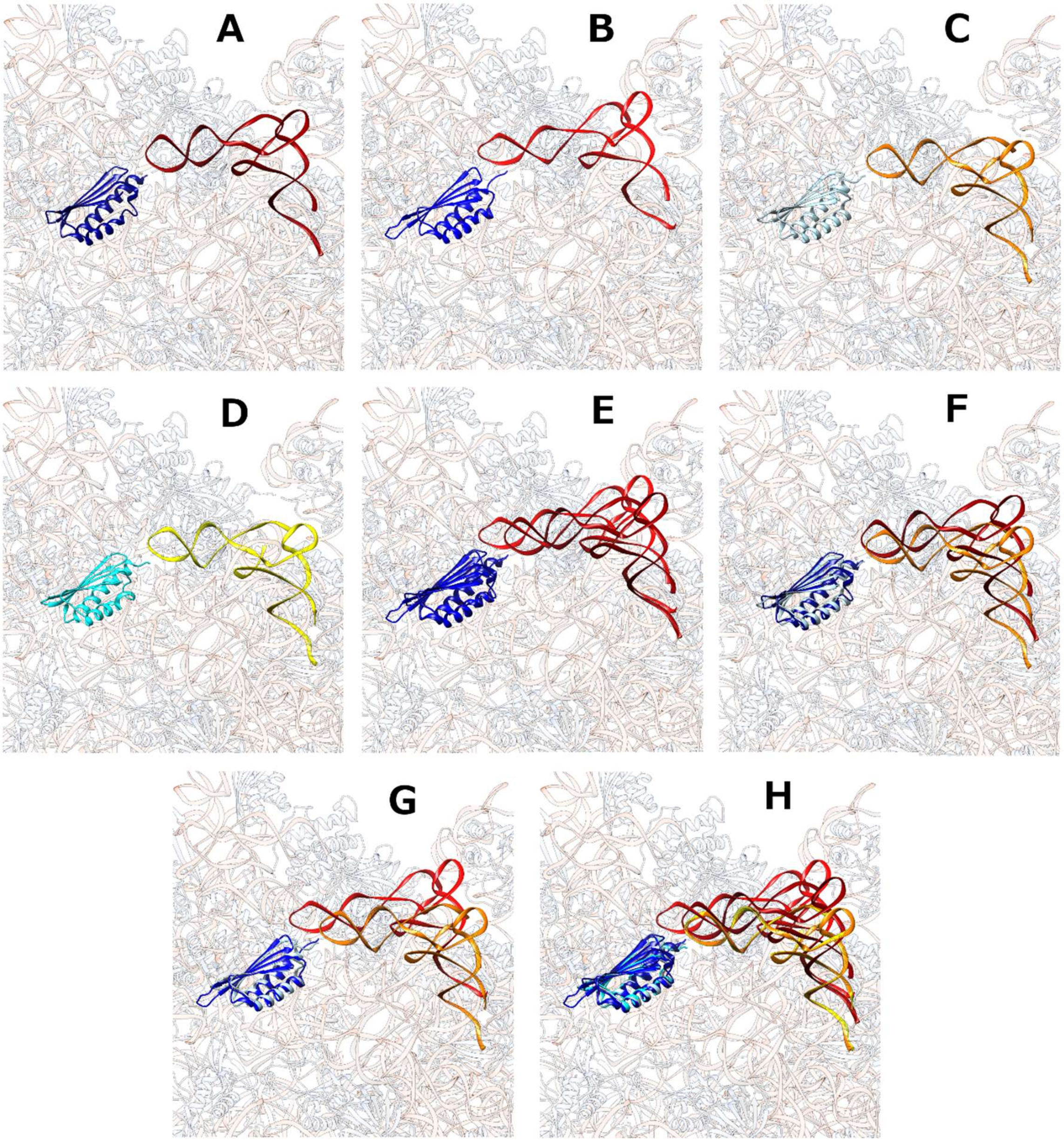


**Figure S2. Variability in relative orientation of HPF protein and E-tRNA bound to 70S ribosomes. (A)** *Bbu* (this study); **(B)** *Msm* (PDB ID: 5ZEP); **(C)** *Eco* (PDB ID: 6H4N); **(D)** *Eco* (PDB ID: 6Y69); **(E)** Overlay of (A) and (B); **(F)** Overlay of (A) and (C); **(G)** Overlay of (B) and (C); **(H)** Overlay of (A)-(D). Background transparent structure is the *Bbu* 70S ribosome.

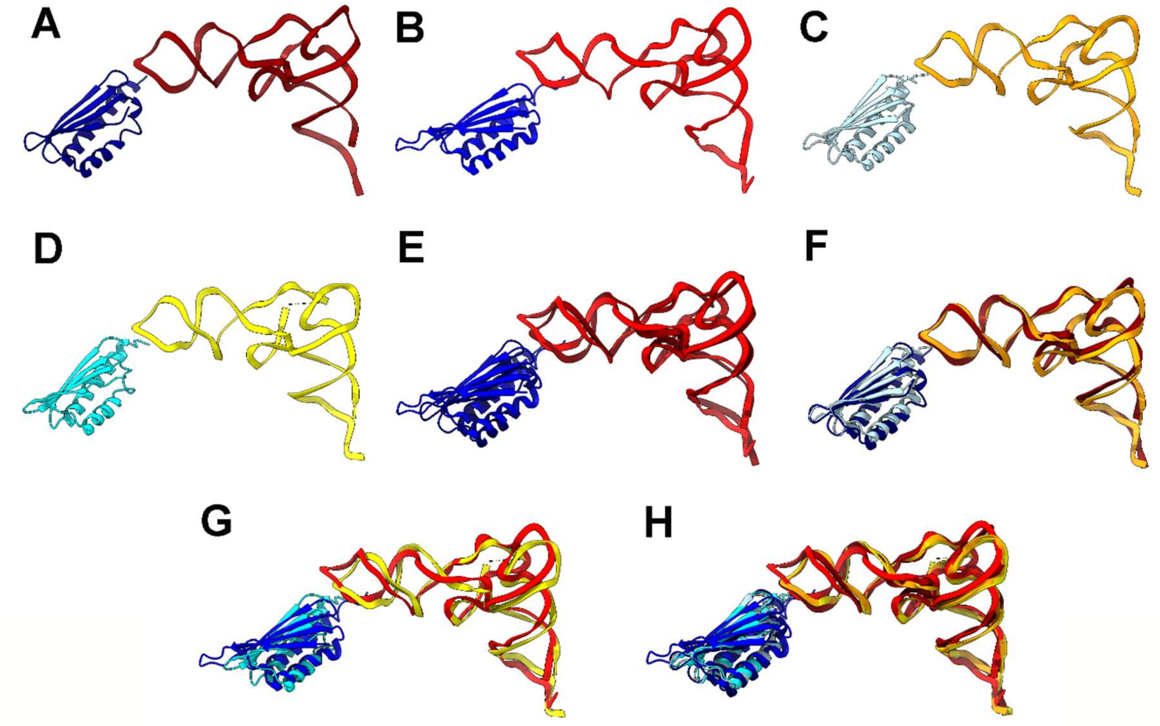


**Figure S3. Adjustment in positioning for both HPF and E-tRNA within their binding sites reduces their overall variability with respect to the 70S ribosome. (A)** *Bbu* (this study); **(B)** *Msm* (PDB ID: 5ZEP); **(C)** *Eco* (PDB ID: 6H4N); **(D)** *Eco* (PDB ID: 6Y69); **(E)** Overlay of (A) and (B); **(F)** Overlay of (A) and (C); **(G)** Overlay of (B) and (C); **(H)** Overlay of (A)-(D). All structures overlaid using the 16S ribosomal RNA.

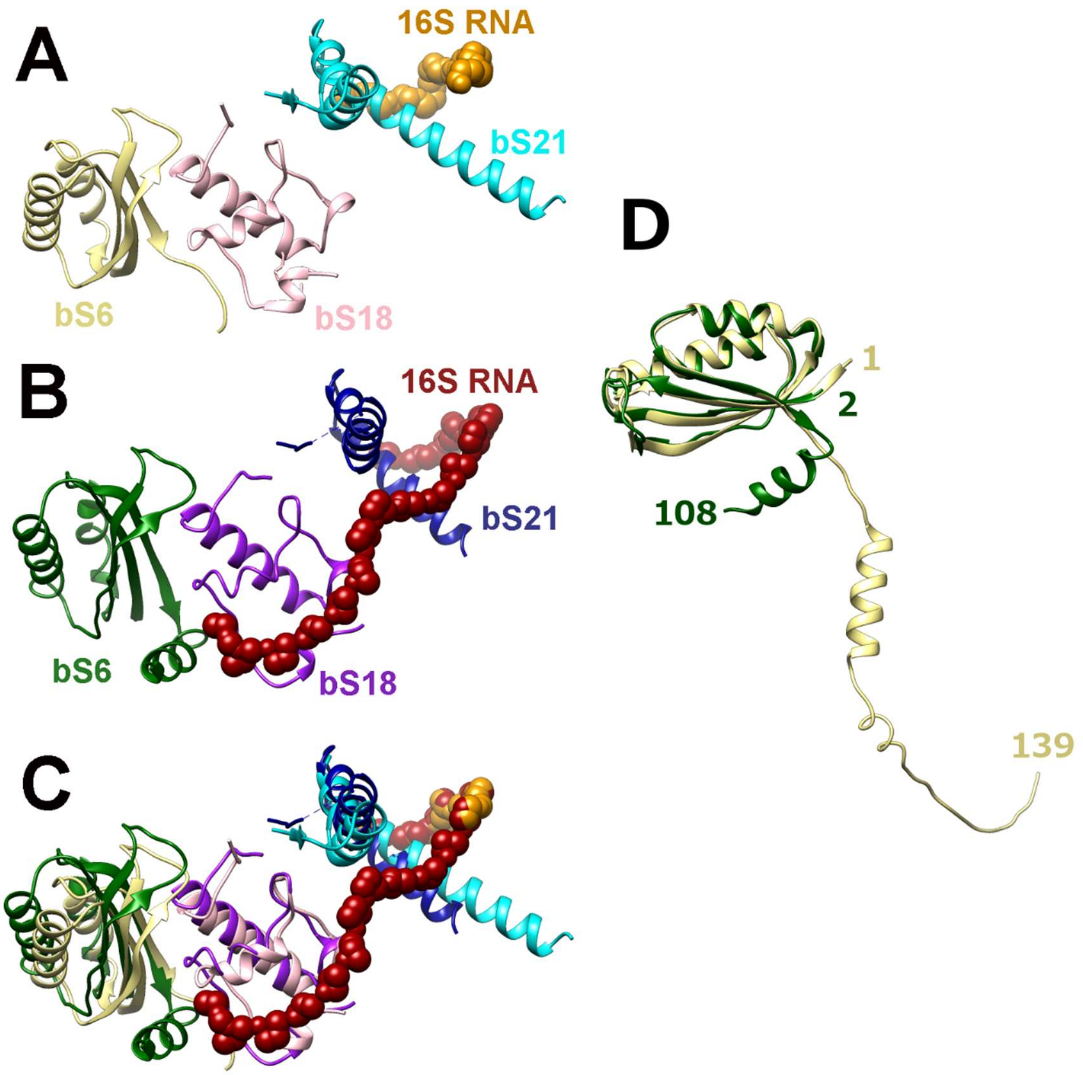


**Figure S4. A binding pocket formed by bS6, bS18 and bS21 implicated in Anti-Shine Dalgarno (ASD) sequence sequestration.** **(A)** The binding pocket in *Bbu* (this study). **(B)** The binding pocket in *Fjo* with the sequestered ASD (PDB ID: 7JIL[5])**. (C)** Overlay of the *Bbu* and *Fjo* binding pockets. **(D)** Overlay of experimentally determined *Fjo* bS6 structure (green) and full-length *Bbu* bS6 Alphafold-predicted structure (yellow) showing likely presence of ASD interacting helix in *Bbu*. For *Bbu*, the 16S RNA minimal backbone -[O5'-C5'-C4'-C3'-O3'-P]- is shown in orange spheres, bS6 is in yellow, bs18 is in pink, and bS21 is in cyan. For *Fjo*, the 16S RNA minimal backbone is shown in red spheres, bS6 is in green, bs18 is in purple, and bS21 is in blue.

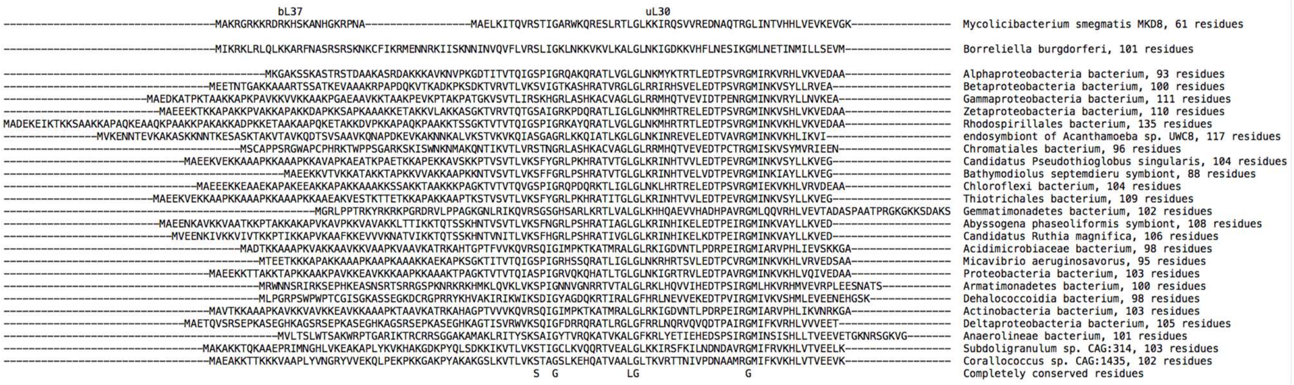


**Figure S5. Long uL30 proteins identified in other bacterial species.** *Msm* ul30 and bL37 sequences are shown on top and the *Bbu* uL30 sequence is shown second for reference.

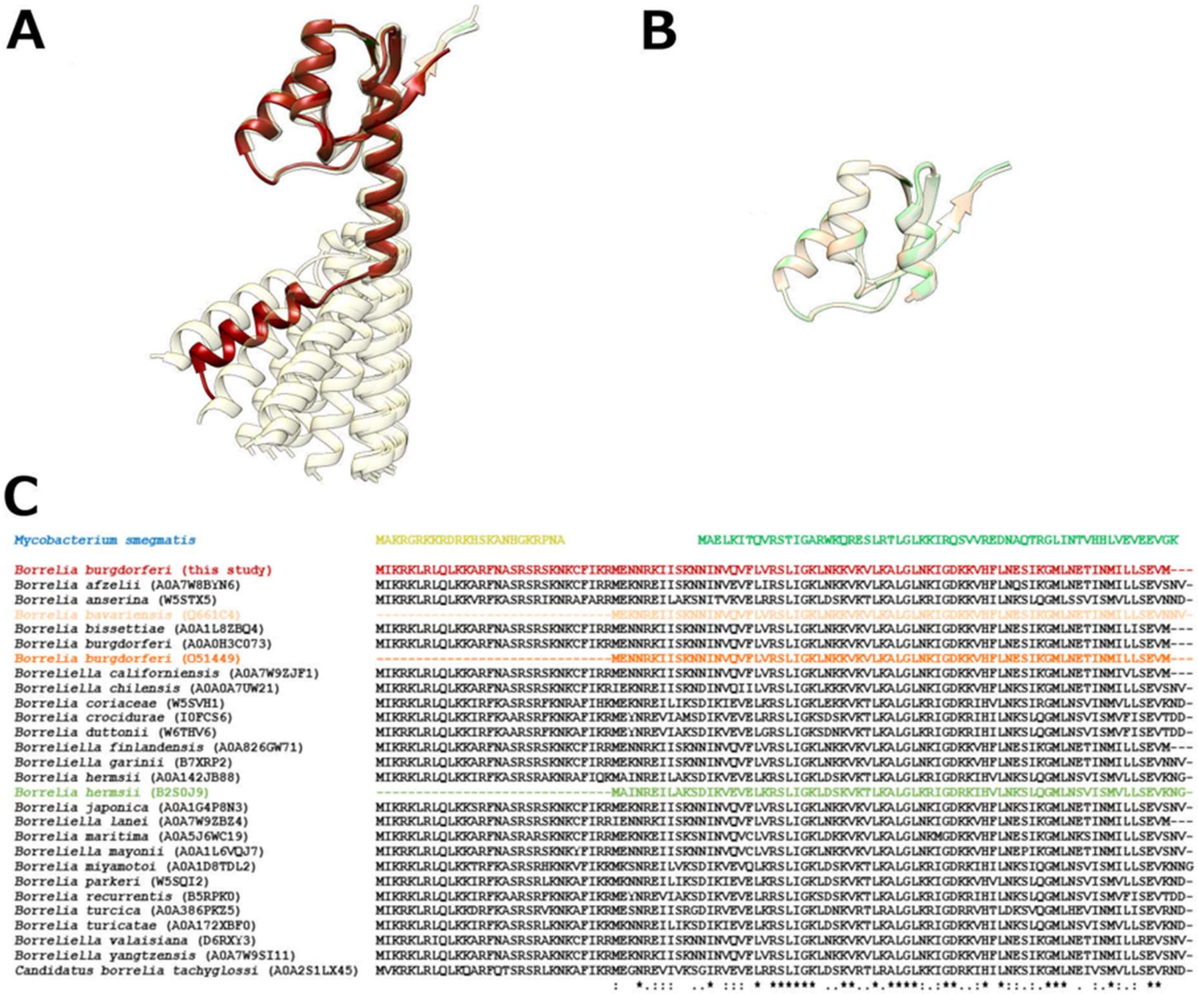


**Figure S6. Two types of longer uL30 proteins in *Borreliella* species. (A)** Alphafold structure predictions for longer *Borreliella* species uL30 proteins, *Bbu* uL30 is shown in dark red; **(B)** Alphafold structure predictions for three shorter *Borreliella* species uL30 proteins; **(C)** Sequence alignment of *Borreliella* species uL30 proteins shown in A and B with *Msm* uL30 and bL37 also aligned for reference. Colors are consistent between models shown in **A** and **B** and sequences shown in **C** except the longer uL30 models shown in khaki color in **A** are shown in black text for their sequence in **C**.

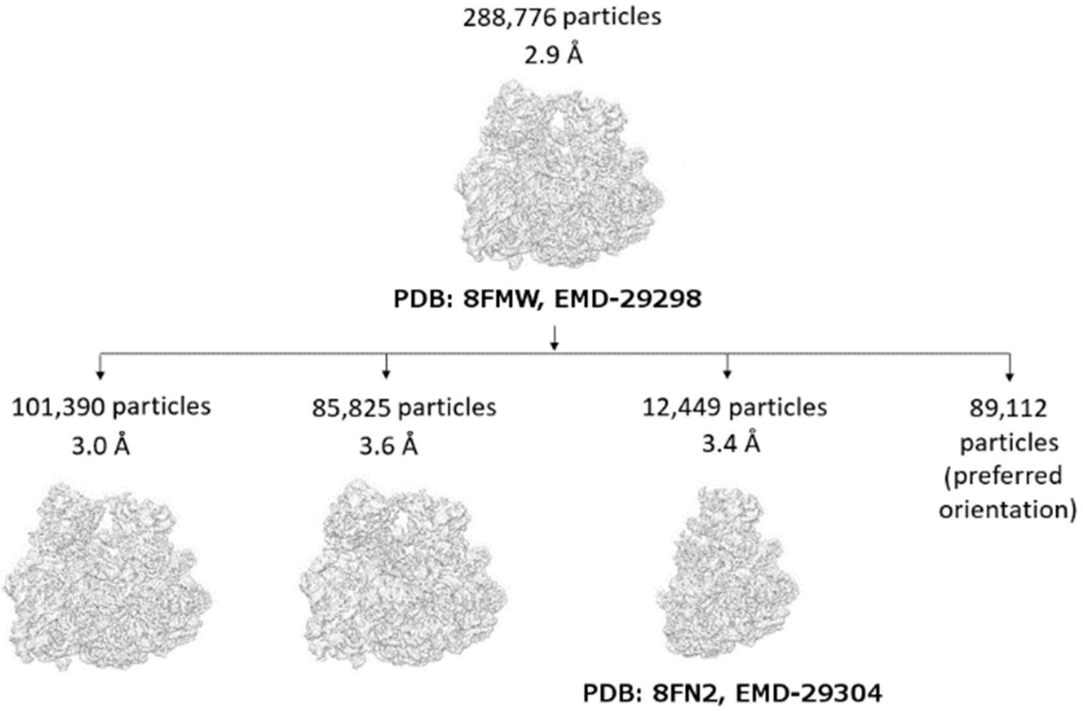


**Figure S7. The 3D classification flowchart of cryo-EM particles for the *Bbu* ribosome.** The classes with 101,390 particles and 85,825 particles did not show any appreciable difference with each other or for subclasses generated from them. The PDB IDs and EMD IDs for the two volumes and corresponding models deposited are shown. All densities are shown at a common threshold value of 0.18.

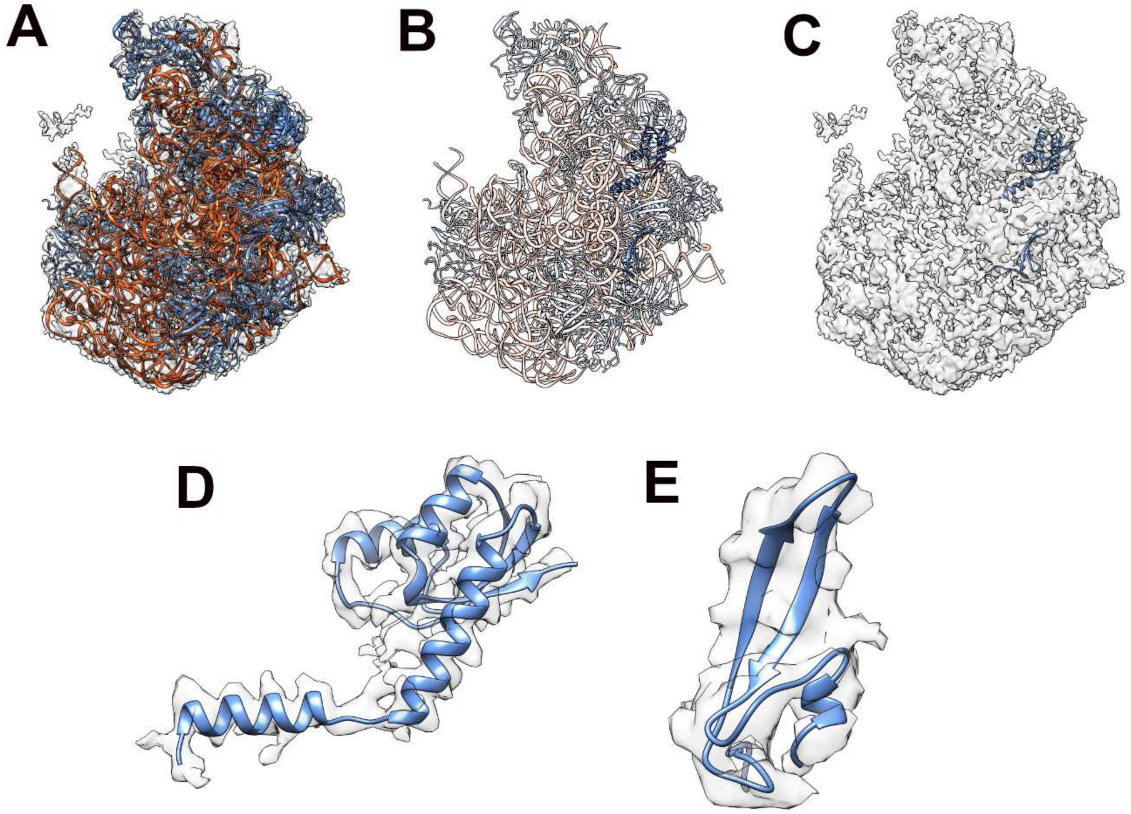


**Figure S8. Large subunit 50S structure for *Bbu* showing that its distinct ribosomal protein components are present in the isolated large subunit and are not specific to the 70S assembly. (A)** The *Bbu* 50S density at 3.4 Å resolution in transparent khaki with its fitted model in opaque orange-red for RNA and in opaque blue for proteins; **(B)** The *Bbu* 50S model in transparent depiction except for uL30 and bL38 proteins shown in opaque blue; **(C)** The *Bbu* 50S density in transparent khaki with uL30 and bL38 proteins shown in opaque blue; **(D)** The model and excised 50S density for uL30; **(E)** The model and excised 50S density for bL38.

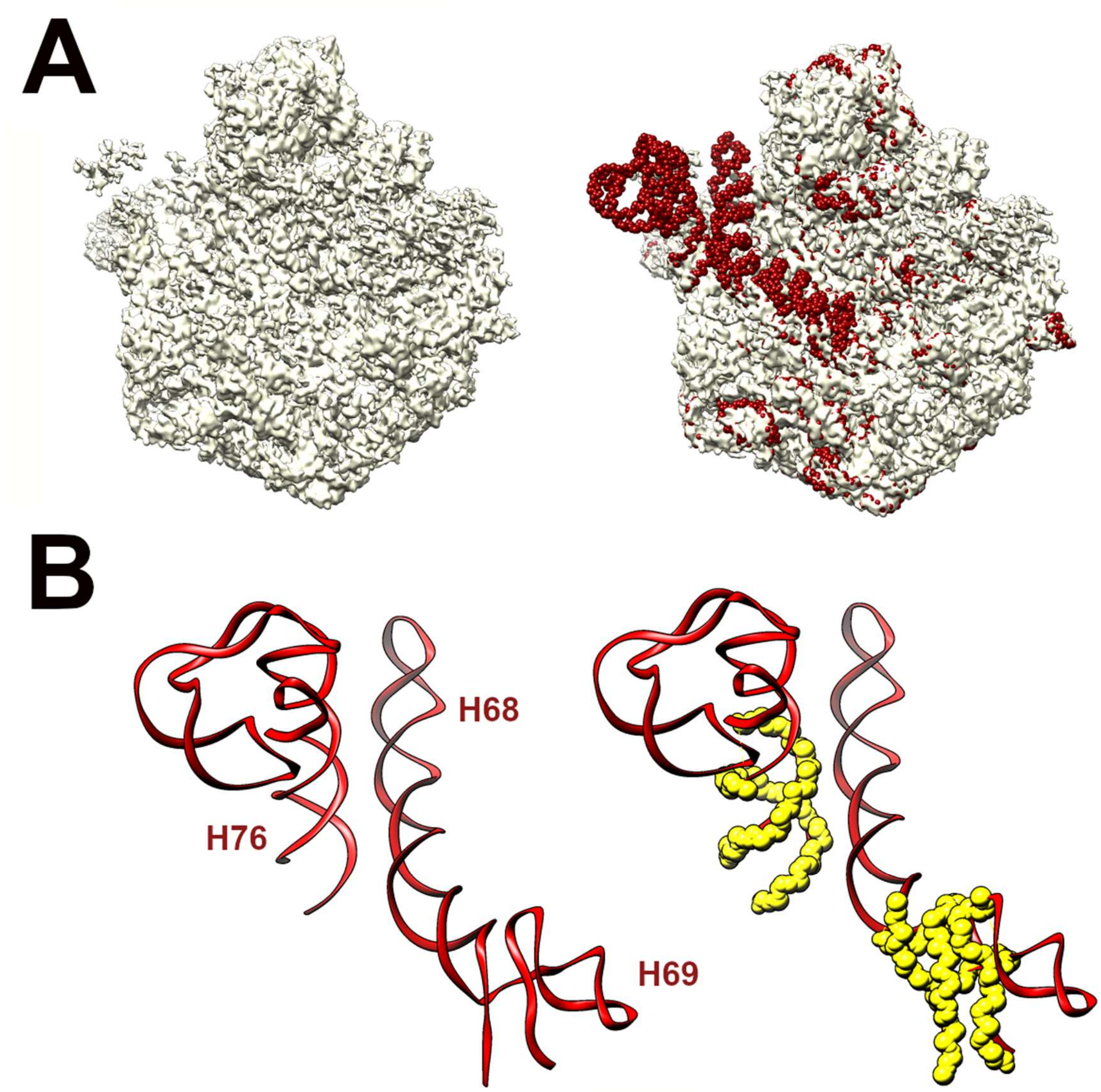


**Figure S9. Disordering of 23S rRNA helices H68 and H69 in the *Bbu* large subunit 50S density. (A)** The *Bbu* 50S density at 3.4 Å resolution in transparent khaki shown at a threshold of 0.15 alone (left) and with the fitted 70S structure 23S rRNA model with backbone atoms shown in red spheres (right) indicating disorder in specific 23S rRNA regions; **(B)** The specific 70S model *Bbu* 23S rRNA regions shown in red ribbons (left) and with the modeled backbone atoms in the 50S 23S rRNA structure overlaid as yellow spheres (right). The helical regions not obscured by the yellow spheres are mostly disordered in the *Bbu* 50S density, suggesting displacement or internal conformation change in 23S RNA helices H68 and H69.

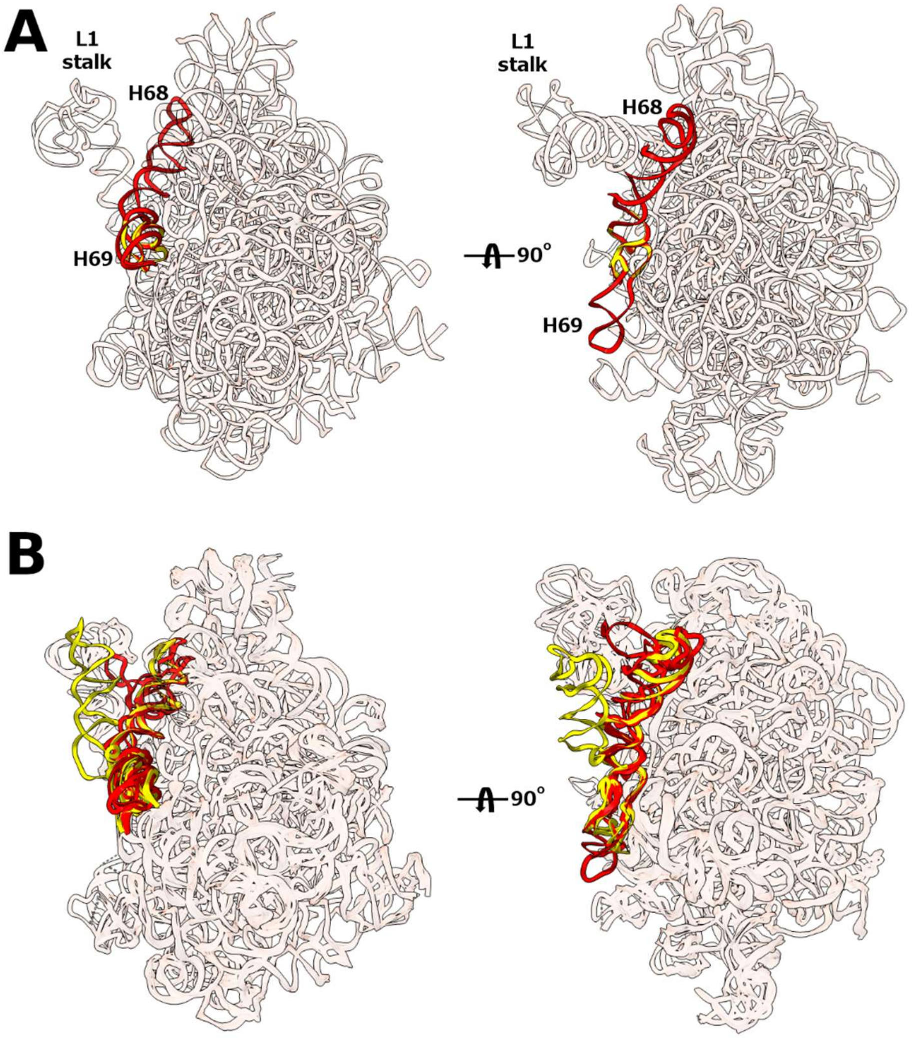


**Figure S10. Comparison of 23S RNA helix 68 (H68) and helix 69 (H69) in 70S and 50S in (A) *Bbu* and (B) *Staphylococcus aureus* (*Sau*).** In panel B, the overlaid *Sau* structures for its 50S and 70S assemblies are shown for 3 durations of incubation at 37°C (in the format: PDB ID for 50S, PDB ID for 70S) as follows: 0 min (6HMA[43], 5TCU[44]), 30 min (7ASM[43], 7ASO[43]), 50 min (7ASN[43], 7ASP[43]). H68 and H69 in the 50S and 70S structures are shown in yellow and red, respectively.

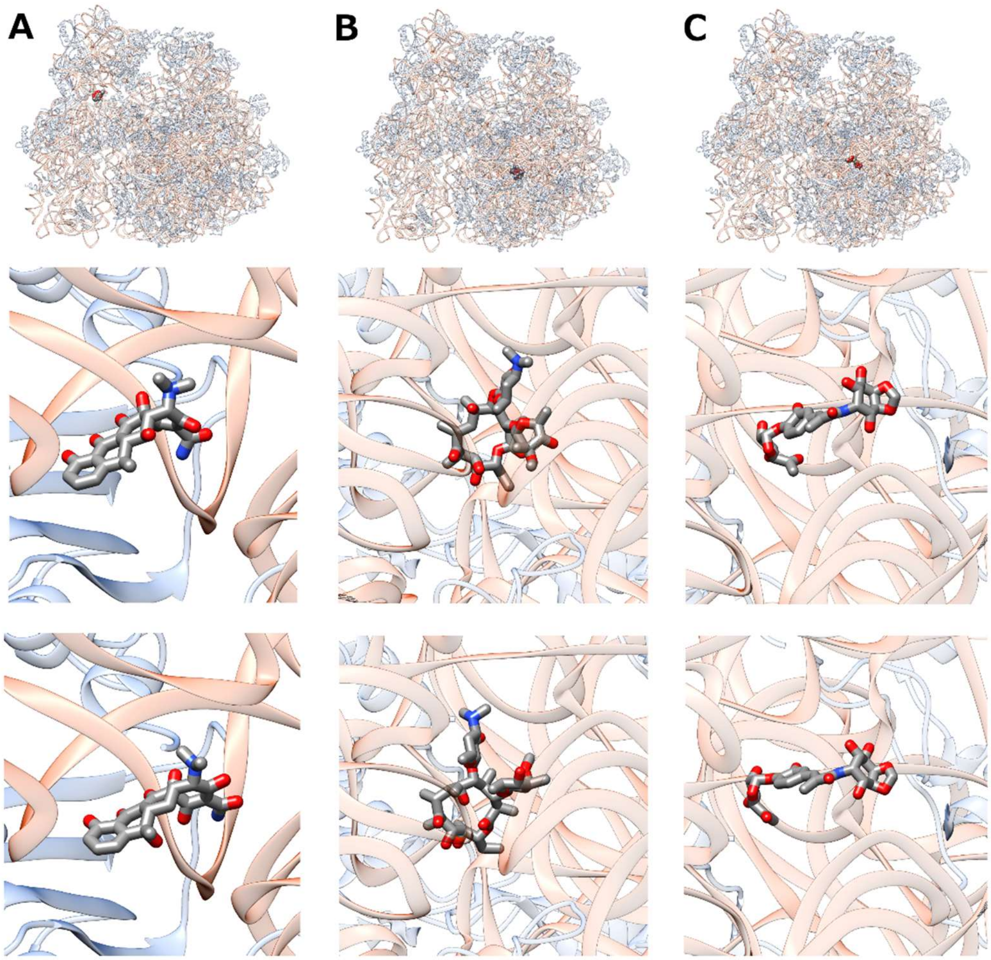


**Figure S11. Antibiotic binding to the *Bbu* 70S ribosome predicted using docking and structural analogy. (A)** Doxycycline docked in the *Bbu* 70S ribosome (top), zoomed in view of docked doxycycline (middle), zoomed in view of tetracycline obtained by structural analogy (bottom, PDB ID: 5J5B[27]); **(B)** Erythromycin docked in the *Bbu* 70S ribosome (top), zoomed in view of docked erythromycin (middle), zoomed in view of erythromycin obtained by structural analogy (bottom, PDB ID: 6S0Z[12]); **(C)** Hygromycin A docked in the *Bbu* 70S ribosome (top), zoomed in view of docked hygromycin A (middle), zoomed in view of hygromycin A obtained by structural analogy (bottom, PDB ID: 5DM7[42]). Docking performed using Quickvina[45] with a box centered around the expected binding site, exhaustiveness parameter set to 32 and number of modes set to 100. Predicted Autodock Vina[46] binding free energies for the *Bbu* 70S ribosome docked positions of the antibiotics shown are as follows: doxycycline -5.4 kcal/mol, erythromycin -6.2 kcal/mol, hygromycin A -7.3 kcal/mol.

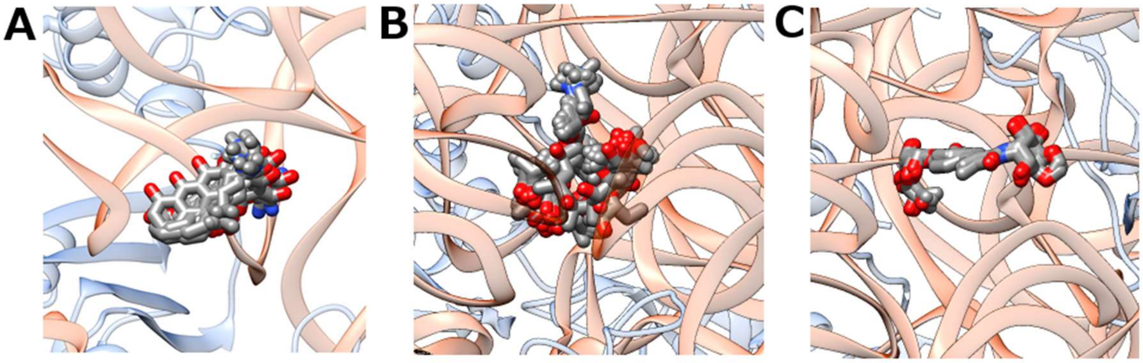

**Figure S12. Multiplicity in antibiotic binding conformations predicted using structural analogy. (A)** Tetracycline in the small subunit decoding center; **(B)** Erythromycin near the peptidyl transferase center (PTC); **(C)** Hygromycin A near the PTC. The antibiotic structures are shown within the *Bbu* 70S ribosome. Structural analogy is obtained by a coarse alignment of the structures with ribosomal subunit RNA and then a finer alignment with neighboring residues within 10 Å of predicted bound antibiotic. Details for structures used in these overlays are in Table S4.

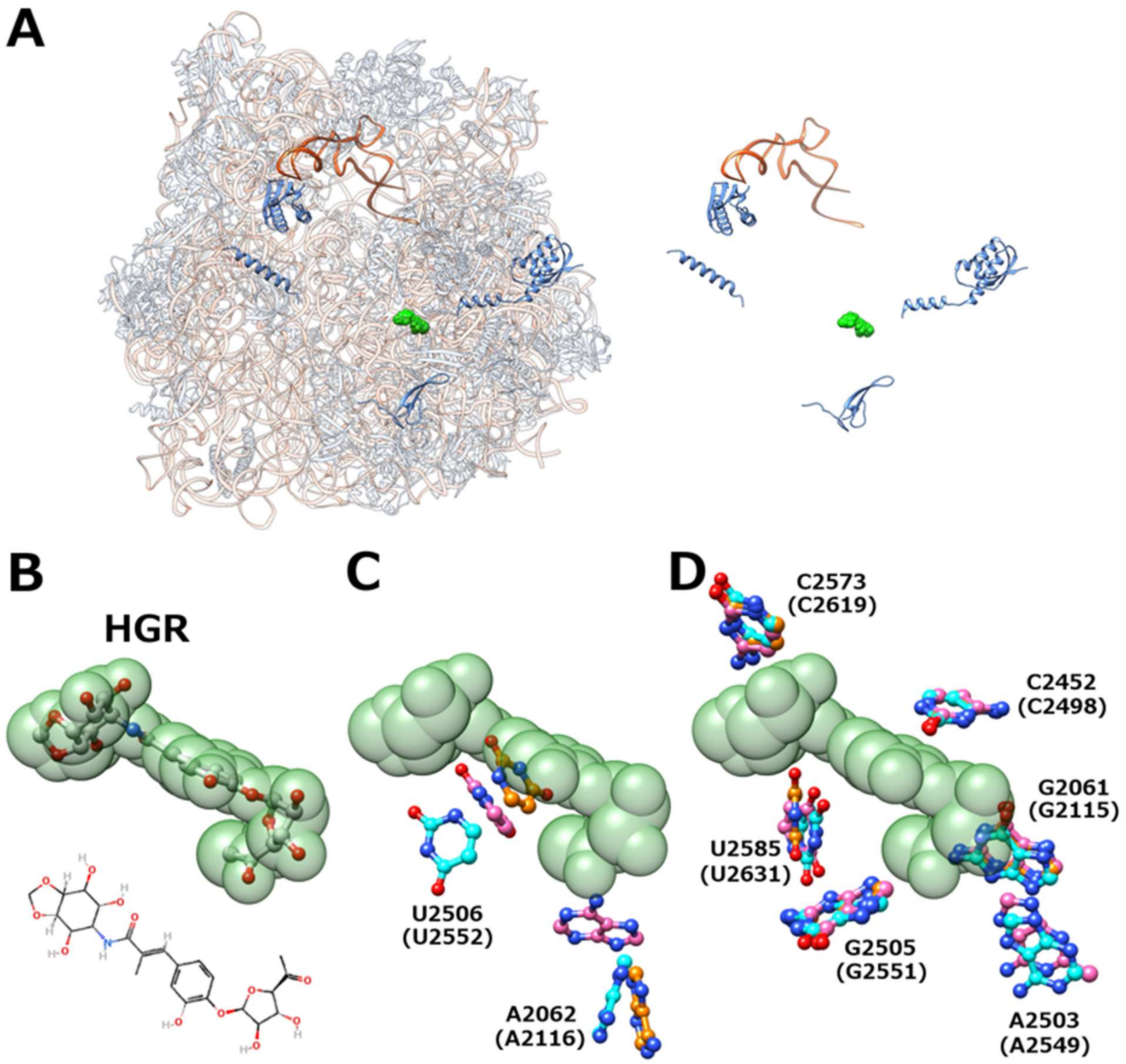


**Figure S13. Comparison of hygromycin A (HGR) binding pocket in *Bbu* and *Tth* ribosomes with and without bound HGR suggests that the empty *Bbu* HGR pocket may be more open for HGR accommodation. (A)** Predicted HGR (green spheres) structure bound in the *Bbu* 50S subunit (left) with its distance from distinct proteins (blue ribbons) indicated (right); **(B)** HGR structure with transparent green spheres overlaid on its atoms (top) and its chemical structure (bottom); **(C)** HGR pocket 23S ribosomal RNA residues showing substantial variability between *Bbu* (this study), *Tth* (PDB ID: 4Y4O[11]), and *Tth* with HGR bound (PDB ID: 5DOX[41]) structures with changes in *Bbu* seemingly making more space for accommodation of HGR; (D) HGR pocket 23S ribosomal RNA residues showing substantial overlap between *Bbu*, *Tth*, and *Tth* HGR-bound structures. Base carbon atoms in 23S ribosomal RNA shown in cyan for *Bbu*, orange for *Tth*, and pink for *Tth* HGR-bound structures. *Bbu* numbering shown for 23S ribosomal RNA residues with *Tth* numbering in parentheses.

**Supplementary movie captions:**

**Supplementary Movie 1:** Movie showing features of the hibernating *Borrelia burgdorferi* (*Bbu*) 70S ribosome structure using a 360° rotation of the soft-lit cryo-EM density of the hibernating *Bbu* 70S ribosome in silver, a second 360° rotation with the cryo-EM density made transparent, a third 360° rotation with the RNA shown in orange-red ribbons, a fourth 360° rotation with the proteins also shown in cornflower-blue ribbons, and a fifth 360° rotation with only the following notable components shown in ribbons: E-tRNA in orange-red, bS22 in cyan, bL38 in green, uL30 in dark-green, and bbHPF in cornflower-blue.

**Supplementary Movie 2:** Movie showing relative motions between hibernation promotion factor (HPF) proteins and E-tRNA using three repetitions of transitions between HPF protein and E-tRNA relative orientations found in four hibernating ribosome structures in *Mycobacterium smegmatis* (*Msm*), *Escherichia coli* (*Eco*) and *Bbu*. For *Bbu*, protein is shown in cornflower-blue ribbons and E-tRNA in dark-red ribbons. For the other organisms, protein is shown in cyan and RNA in orange.

**Supplementary Movie 3:** Movie showing analogy between the uL30 protein in *Bbu* with large subunit ribosomal proteins in other organisms using a 360° rotation of the *Bbu* uL30 protein in cornflower-blue ribbons, a 360° rotation of the *Msm* uL30 protein in green ribbons and the bL37 protein in yellow ribbons, a 360° rotation of the *Homo sapiens* (*Hsa*) uL30 protein in orange ribbons, a 360° rotation of the *Hsa* mitochondrial uL30m protein in green ribbons and the mL63 protein in yellow ribbons, a 360° rotation of an overlay of the *Bbu* uL30 protein and the *Msm* uL30 and bL37 proteins, a 360° rotation of an overlay of the *Bbu* uL30 protein and the *Hsa* uL30 protein, and a 360° rotation of an overlay of the *Bbu* uL30 protein and the *Hsa* mitochondrial uL30m protein and the mL63 protein.

**Supplementary Movie 4:** Movie showing 23S RNA H68 and H69 disorder or motions using three repetitions of transitions between oriented 23S RNA structures found in eight ribosome structures from two bacteria: *Bbu* and *Staphylococcus aureus* (*Sau*), followed by their overlay. The structures are: *Bbu* 50S (PDB ID: 8FN2), *Bbu* 70S (PDB ID: 8FMW), *Sau* 50S with 0 minutes incubation at 37 °C (PDB ID: 6HMA), *Sau* 50S with 30 minutes incubation at 37 °C (PDB ID: 7ASM), *Sau* 50S with 50 minutes incubation at 37 °C (PDB ID: 7ASN), *Sau* 70S with 0 minutes incubation at 37 °C (PDB ID: 5TCU), *Sau* 70S with 30 minutes incubation at 37 °C (PDB ID: 7ASO), *Sau* 70S with 50 minutes incubation at 37 °C (PDB ID: 7ASP). For *Bbu*, 23S RNA residues 1888-2001 are shown in green, for *Sau* corresponding residues 1861-1974 are shown in yellow.

### References.


1 Hentschel, J. *et al.* The complete structure of the Mycobacterium smegmatis 70S ribosome. *Cell reports* **20**, 149-160 (2017).

2 Li, Z. *et al.* Cryo-EM structure of Mycobacterium smegmatis ribosome reveals two unidentified ribosomal proteins close to the functional centers. *Protein & cell* **9**, 384-388 (2018).

3 Li, Y. *et al.* Zinc depletion induces ribosome hibernation in mycobacteria. *Proc Natl Acad Sci U S A* **115**, 8191-8196, doi:10.1073/pnas.1804555115 (2018).

4 Mishra, S., Ahmed, T., Tyagi, A., Shi, J. & Bhushan, S. Structures of Mycobacterium smegmatis 70S ribosomes in complex with HPF, tmRNA, and P-tRNA. *Scientific reports* **8**, 1-12 (2018).

5 Jha, V. *et al.* Structural basis of sequestration of the anti-Shine-Dalgarno sequence in the Bacteroidetes ribosome. *Nucleic acids research* **49**, 547-567 (2021).

6 Gertz, E. M., Yu, Y.-K., Agarwala, R., Schäffer, A. A. & Altschul, S. F. Composition-based statistics and translated nucleotide searches: improving the TBLASTN module of BLAST. *BMC biology* **4**, 1-14 (2006).

7 Cao, X. & Slavoff, S. A. Non-AUG start codons: Expanding and regulating the small and alternative ORFeome. *Experimental cell research* **391**, 111973 (2020).

8 Villegas, A. & Kropinski, A. M. An analysis of initiation codon utilization in the Domain Bacteria–concerns about the quality of bacterial genome annotation. *Microbiology* **154**, 2559-2661 (2008).

9 Thompson, J. D., Gibson, T. J. & Higgins, D. G. Multiple sequence alignment using ClustalW and ClustalX. *Current protocols in bioinformatics*, 2.3. 1-2.3. 22 (2003).

10 De Bari, H. & Berry, E. A. Structure of Vibrio cholerae ribosome hibernation promoting factor. *Acta Crystallographica Section F: Structural Biology and Crystallization Communications* **69**, 228-236 (2013).

11 Polikanov, Y. S., Melnikov, S. V., Söll, D. & Steitz, T. A. Structural insights into the role of rRNA modifications in protein synthesis and ribosome assembly. *Nature structural & molecular biology* **22**, 342-344 (2015).

12 Halfon, Y. *et al.* Exit tunnel modulation as resistance mechanism of S. aureus erythromycin resistant mutant. *Scientific reports* **9**, 1-8 (2019).

13 Syroegin, E. A. *et al.* Structural basis for the context-specific action of the classic peptidyl transferase inhibitor chloramphenicol. *Nature Structural & Molecular Biology* **29**, 152-161 (2022).

14 Franklin, M. C. *et al.* Structural genomics for drug design against the pathogen Coxiella burnetii. *Proteins: Structure, Function, and Bioinformatics* **83**, 2124-2136 (2015).

15 Tereshchenkov, A. G. *et al.* Binding and action of amino acid analogs of chloramphenicol upon the bacterial ribosome. *Journal of molecular biology* **430**, 842-852 (2018).

16 Svetlov, M. S. *et al.* Structure of Erm-modified 70S ribosome reveals the mechanism of macrolide resistance. *Nature chemical biology* **17**, 412-420 (2021).

17 Polikanov, Y. S., Blaha, G. M. & Steitz, T. A. How hibernation factors RMF, HPF, and YfiA turn off protein synthesis. *Science* **336**, 915-918 (2012).

18 Seefeldt, A. C. *et al.* Structure of the mammalian antimicrobial peptide Bac7 (1–16) bound within the exit tunnel of a bacterial ribosome. *Nucleic acids research* **44**, 2429-2438 (2016).

19 Chen, C.-W. *et al.* Binding and action of triphenylphosphonium analog of chloramphenicol upon the bacterial ribosome. *Antibiotics* **10**, 390 (2021).

20 Matzov, D. *et al.* The cryo-EM structure of hibernating 100S ribosome dimer from pathogenic Staphylococcus aureus. *Nature communications* **8**, 1-7 (2017).

21 Osterman, I. A. *et al.* Tetracenomycin X inhibits translation by binding within the ribosomal exit tunnel. *Nature Chemical Biology* **16**, 1071-1077 (2020).

22 Zhang, Z., Morgan, C. E., Bonomo, R. A. & Yu, E. W. Cryo-EM determination of Eravacycline-Bound structures of the Ribosome and the multidrug efflux pump AdeJ of Acinetobacter baumannii. *MBio* **12**, e01031-01021 (2021).

23 Beckert, B. *et al.* Structure of a hibernating 100S ribosome reveals an inactive conformation of the ribosomal protein S1. *Nature microbiology* **3**, 1115-1121 (2018).

24 Mardirossian, M. *et al.* The dolphin proline-rich antimicrobial peptide Tur1A inhibits protein synthesis by targeting the bacterial ribosome. *Cell chemical biology* **25**, 530-539. e537 (2018).

25 Flygaard, R. K., Boegholm, N., Yusupov, M. & Jenner, L. B. Cryo-EM structure of the hibernating Thermus thermophilus 100S ribosome reveals a protein-mediated dimerization mechanism. *Nature communications* **9**, 1-12 (2018).

26 Li, Y. *et al.* Zinc depletion induces ribosome hibernation in mycobacteria. *Proceedings of the National Academy of Sciences* **115**, 8191-8196 (2018).

27 Cocozaki, A. I. *et al.* Resistance mutations generate divergent antibiotic susceptibility profiles against translation inhibitors. *Proceedings of the National Academy of Sciences* **113**, 8188-8193 (2016).

28 Jenner, L. *et al.* Structural basis for potent inhibitory activity of the antibiotic tigecycline during protein synthesis. *Proceedings of the National Academy of Sciences* **110**, 3812-3816 (2013).

29 Brodersen, D. E. *et al.* The structural basis for the action of the antibiotics tetracycline, pactamycin, and hygromycin B on the 30S ribosomal subunit. *Cell* **103**, 1143-1154 (2000).

30 Pioletti, M. *et al.* Crystal structures of complexes of the small ribosomal subunit with tetracycline, edeine and IF3. *The EMBO journal* **20**, 1829-1839 (2001).

31 Tu, D., Blaha, G., Moore, P. B. & Steitz, T. A. Structures of MLSBK antibiotics bound to mutated large ribosomal subunits provide a structural explanation for resistance. *Cell* **121**, 257-270 (2005).

32 Svetlov, M. S. *et al.* High-resolution crystal structures of ribosome-bound chloramphenicol and erythromycin provide the ultimate basis for their competition. *Rna* **25**, 600-606 (2019).

33 Albers, S. *et al.* Repurposing tRNAs for nonsense suppression. *Nature Communications* **12**, 3850 (2021).

34 Beckert, B. *et al.* Structural and mechanistic basis for translation inhibition by macrolide and ketolide antibiotics. *Nature Communications* **12**, 4466 (2021).

35 Bulkley, D., Innis, C. A., Blaha, G. & Steitz, T. A. Revisiting the structures of several antibiotics bound to the bacterial ribosome. *Proceedings of the National Academy of Sciences* **107**, 17158-17163 (2010).

36 Wekselman, I. *et al.* The ribosomal protein uL22 modulates the shape of the protein exit tunnel. *Structure* **25**, 1233-1241. e1233 (2017).

37 Schlünzen, F. *et al.* Structural basis for the interaction of antibiotics with the peptidyl transferase centre in eubacteria. *Nature* **413**, 814-821 (2001).

38 Arenz, S. *et al.* A combined cryo-EM and molecular dynamics approach reveals the mechanism of ErmBL-mediated translation arrest. *Nature communications* **7**, 12026 (2016).

39 Arenz, S. *et al.* Drug sensing by the ribosome induces translational arrest via active site perturbation. *Molecular cell* **56**, 446-452 (2014).

40 Arenz, S. *et al.* Molecular basis for erythromycin-dependent ribosome stalling during translation of the ErmBL leader peptide. *Nature communications* **5**, 3501 (2014).

41 Polikanov, Y. S. *et al.* Distinct tRNA accommodation intermediates observed on the ribosome with the antibiotics hygromycin A and A201A. *Molecular cell* **58**, 832-844 (2015).
42 Kaminishi, T. *et al.* Crystallographic characterization of the ribosomal binding site and molecular mechanism of action of Hygromycin A. *Nucleic acids research* **43**, 10015-10025 (2015).
43 Cimicata, G. *et al.* Structural Studies Reveal the Role of Helix 68 in the Elongation Step of Protein Biosynthesis. *Mbio* **13**, e00306-00322 (2022).
44 Belousoff, M. J. *et al.* Structural basis for linezolid binding site rearrangement in the Staphylococcus aureus ribosome. *MBio* **8**, e00395-00317 (2017).
45 Alhossary, A., Handoko, S. D., Mu, Y. & Kwoh, C.-K. Fast, accurate, and reliable molecular docking with QuickVina 2. *Bioinformatics* **31**, 2214-2216 (2015).
46 Trott, O. & Olson, A. J. AutoDock Vina: improving the speed and accuracy of docking with a new scoring function, efficient optimization, and multithreading. *Journal of computational chemistry* **31**, 455-461 (2010).
